# Dissecting conformational rearrangements and allosteric modulation in metabotropic glutamate receptor activation

**DOI:** 10.1101/2022.01.07.474531

**Authors:** Nathalie Lecat-Guillet, Robert B. Quast, Hongkang Liu, Thor C. Møller, Xavier Rovira, Stéphanie Soldevila, Laurent Lamarque, Eric Trinquet, Jianfeng Liu, Jean-Philippe Pin, Philippe Rondard, Emmanuel Margeat

## Abstract

Selective allosteric modulators bear great potential to fine-tune neurotransmitter-induced brain receptor responses. Promising targets are metabotropic glutamate (mGlu) receptors, which are associated to different brain diseases. These multidomain class C GPCRs experience concerted structural rearrangements and rely on allosteric modulation of agonist action to be fully activated. Here we establish live cell compatible fluorescence labeling of mGlu2 by click chemistry through genetic code expansion. Using lanthanide resonance energy transfer, we establish multiple FRET sensors to monitor ligand effects on conformational changes in mGlu2 extracellular domain and subsequently dissect the underlying conformational states by smFRET. Using three distinct FRET sensors, we demonstrate that mGlu activation relies on a ligand-induced sampling of three conformational states. Orthosteric agonists act by promoting the closure of the mGlu2 ligand binding domains, leading to an equilibrium between an inactive intermediate and the active state. Allosteric modulator further push this equilibrium toward the active state, promoting and stabilizing the relative reorientation of the mGlu protomers. These results underline the complex and dynamic nature of such type of neuroreceptors, pointing out that ligands fine-tune activation by differentially acting on the equilibria between multiple states.

## Introduction

The family of metabotropic glutamate (mGlu) receptors comprises eight types of neuroreceptors sensing L-glutamate, the major excitatory neurotransmitter in the human brain^1^. Their role in regulating neuronal excitability and synaptic transmission makes them important targets for the treatment of neurological and psychiatric diseases^2, 3^. A major challenge in drug discovery arises from the high conservation of the orthosteric ligand-binding site between different members of the mGlu family^4^. This has directed drug discovery efforts towards small molecular allosteric modulators that target other binding pockets and therefore show higher subtype specificity^5, 6^.

mGlu receptors are dimeric and multidomain proteins belonging to the class C of G protein-coupled receptors (GPCR). They share a similar structure and mechanism of activation with the other class C GPCRs activated by L-amino-acids and cations such as the calcium sensing receptor, the GPRC6A, the umami and sweet taste receptors^7^. In these receptors, orthosteric ligands bind to a cleft within the upper and lower lobes of the Venus flytrap domain (VFT), inducing large conformational rearrangements to activate the receptor followed by recruitment of downstream signaling partners interacting with the cytoplasmic side of the 7 transmembrane-spanning domain (7TM), most notably G proteins **(Figure 1a)**. Early structural studies of isolated VFTs of mGlu receptors have shown that the upper and lower lobes come into closer proximity when activated by agonists, leading to a closure of the two VFTs (open-open “oo” to closed-closed “cc” conformation). This leads to a reorientation of the two adjacent VFT protomers, increasing the distance between the upper lobes, while the lower lobes reorient to approach each other (resting “R” to active “A” conformation) **(Figure 1a)**^8–11^. Such a model highlighting the Roo and the Acc conformations was recently confirmed by cryo-EM structures of nearly full-length receptors^12–15^. They also revealed a non-conserved mechanism with regard to the reorganization of the 7TM helix bundle upon activation, compared to GPCRs from other classes^14, 16^. Surprisingly, some early crystal VFT structures showed agonist-bound mGlu3 VFT dimers, adopting a Rcc conformation with the VFTs being closed but the lower lobes remaining separated^10^, implying that agonists alone may not be sufficient for full VFT reorientation. In accordance with this, we recently used single molecule Förster resonance energy transfer (smFRET) on receptors labeled through N-terminal SNAP-tags to show that the combination of an orthosteric agonist with positive allosteric modulators (such as the small organic molecule BINA or the heterotrimeric G protein) is required for promoting a full shift of the equilibrium toward the active state ^17^. This SNAP sensor only reported on the reorientation of the N-termini being located on the upper lobes of the VFTs (**Figure 1a**). However, fully understanding how ligands act on the initial steps of mGlu receptor activation requires to be able to quantify their effect on the structural dynamics of the receptor at the level of each of its domains, including for example the closure of the VFT, or the relative reorientation of the VFTs, of the transmembrane domains, or of the cysteine-rich domains (CRD) connecting them.

**Figure 1:**
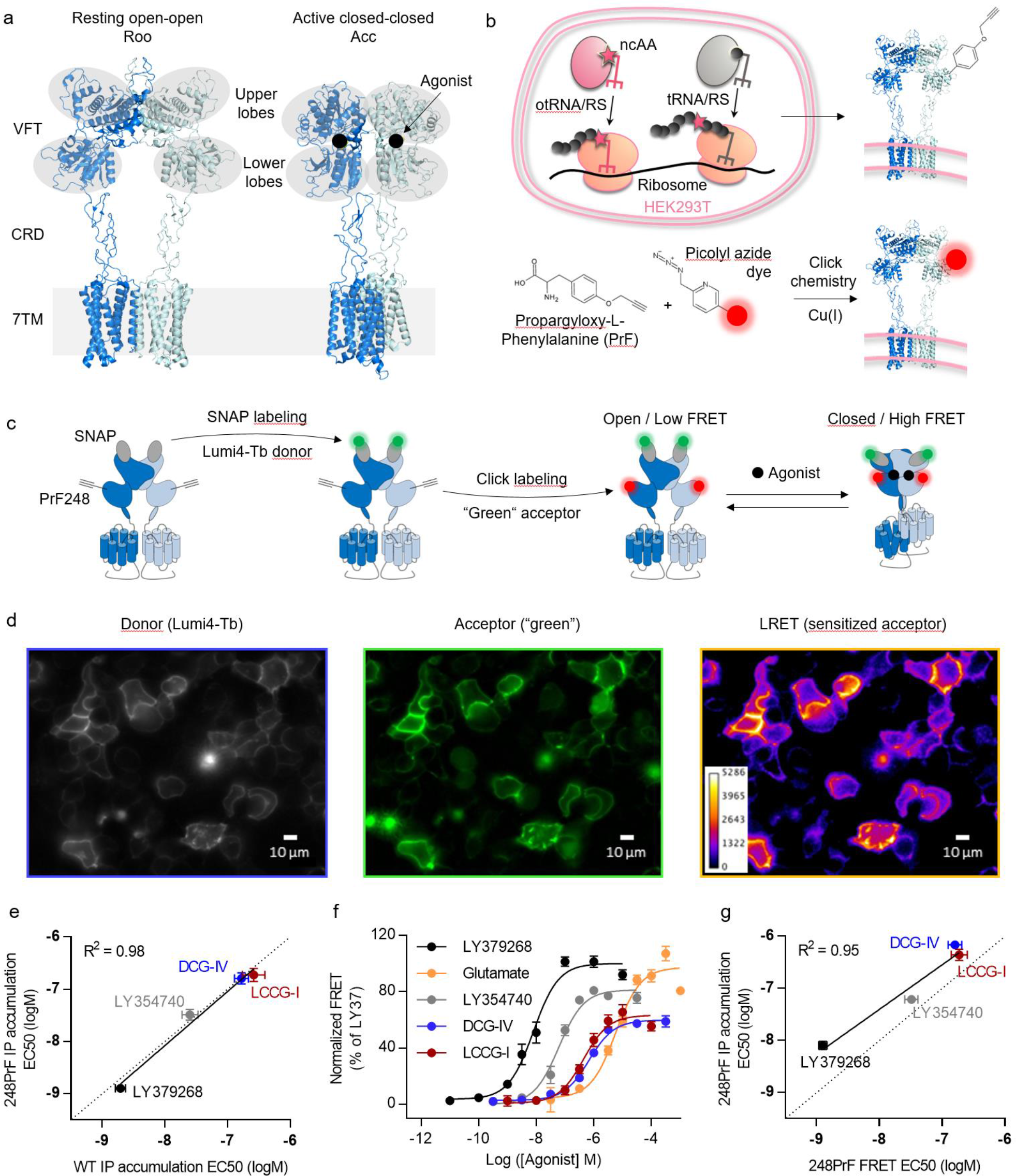
Site-specific fluorescence labeling of mGlu receptors by genetic code expansion and bioorthogonal click chemistry. a) Resting open-open (Roo) and active closed-closed (Acc) structures of mGlu2 (PDB 7EPA, 7E9G). The major domains are annotated including the upper and lower lobes of the Venus flytrap domain (VFT), the cysteine-rich domaine (CRD) and the 7-transmembrane spanning domain (7TM). The upper and lower lobes are highlighted in gray and agonist-binding to the orthosteric site is shown as black spheres. b) Schematic representation of genetic code expansion and bioorthogonal click chemistry. Co-expression of the orthogonal synthetase together with its cognate amber suppressor tRNACUA (otRNA/RS) in the presence of the non-canonical amino acid Propargyloxyphenylalanine (PrF) in HEK293T cells allows the reassignment of a premature Amber codon TAG within the ORF of the receptor gene to code for PrF. This leads to incorporation of PrF at a predefined position in the translated receptor, which can subsequently be selectively reacted with a Picolyl azide (pAz) dye using Copper(I)-catalyzed azide-alkyne cycloaddition (CuAAC). c) Schematic representation of N-terminal SNAP-labeling followed by click labeling of the lower lobe (A248) to obtain the FRET sensor used in panels d-g. d) LRET imaging of mGlu2 labeled with the SNAP-Lumi4-Tb donor at N-terminal SNAP-tags and at position 248 in the lower lobe of the VFT with the green Picolyl azide acceptor at the plasma membrane of live HEK293T cells. e) Comparison of logEC50 values of ligand- induced receptor activation of mGlu2 with incorporated PrF (248PrF) and wildype receptor (WT) in living cells using different ligands based on the accumulation of phosphoinositol monophosphate (IP). f) Dose-response curves showing ligand-induced increase of LRET as a result of VFT closure on mGlu2 receptors. g) Comparison of logEC50 values for different ligands, determined by IP accumulation and LRET. Data in e-g represent the mean of 3-12 measurements +/- SD.

Therefore, we established a strategy to generate a library of site-specific FRET sensors to monitor the individual domain reorientations, focusing on the mGlu2 receptor VFTs. It relies on the incorporation of a reactive non-canonical amino acid (ncAA) by genetic code expansion (**Figure 1b**). In this way FRET-compatible donor and acceptor fluorophores can be installed at desired positions using bioorthogonal labeling strategies^18, 19^. First, we screened for suitable labeling positions based on ncAA incorporation efficiency and functional integrity of modified receptors, followed by the establishment of live cell compatible click chemistry conditions that allow to monitor ligand-induced conformational rearrangements by lanthanide resonance energy transfer (LRET). For three selected FRET sensors, full-length receptors were then detergent-solubilized and analyzed at the single molecule level. By quantifying the effect of orthosteric and allosteric ligands on i) the intrasubunit closure of the VFT, ii) the intersubunit rearrangement of the upper lobes and iii) the intersubunit reduction in lower lobe separation, we derive a three-state model composed of two main conformational equilibria between the Ropen, Rclosed and Aclosed states. On one hand, we show that orthosteric agonists act by promoting the closure of the VFT domain (open to closed equilibrium), and that the fraction of closed VFTs is correlated with their receptor activation efficacies. On the other hand, we show that the positive allosteric modulator (PAM) BINA is mainly involved in promoting the reorientation of the VFT dimer, acting on the R-A equilibrium and thereby leading to maximal activation of the receptor.

## Results

### Live cell fluorescent labeling of ncAAs in mGlu2 by click chemistry

Firstly, we established conditions for the site-specific incorporation and efficient labeling of the ncAA 4- Propargyloxy-L-phenylalanine (PrF), in combination with copper-dependent azide-alkyne cycloaddition (CuAAC) using picolyl azide (pAz) dyes (**Figure 1b**). Therefore, we introduced premature Amber codons (TAG) at 13 different positions within the gene of our SNAP-FLAG-mGlu2 construct throughout the region coding for the VFT (**Figure S1**). We selected the ncAA PrF due to its very good stability under physiological conditions and its capability to be reacted with aryl-azides in the presence of Cu(I) through CuAAC in a fast, highly-selective and bioorthogonal manner^20^. PrF was incorporated in response to the suppression of premature TAG by co-transfection of HEK293T cells with the vector coding for SNAP-mGlu2 together with a bicistronic vector coding for an engineered *E. coli* tyrosly-tRNA synthetase (PrFRS)^21, 22^ and three copies of the engineered *B. stearothermophilus* tyrosyl-tRNACUA. By screening the various positions, we found that substituting Ala248 within the lower lobe of the VFT provided good incorporation yields in a PrF-dependent manner compared to the expression of the wildtype receptor and correspondingly modified receptors maintained signaling activity (**Figure S1** and **S2a**). Therefore, we further focused on this SNAP-mGlu2-PrF248 mutant to optimize the CuAAC labeling conditions.

The CuAAC reaction requires the presence of Cu(I) to proceed efficiently under physiological buffer conditions^24, 25^. Multiple protocols have been described to reduce Cu(II) to Cu(I), stabilize it in solution and minimize its cytotoxicity as well as that of unwanted side products, e.g. reactive oxygen species. With the aim of performing LRET measurements of receptors within the plasma membrane of living cells, we combined two previously described strategies to minimize the required concentration of copper, while maintaining satisfying reaction efficiencies. Firstly, we synthesized pAz derivatives of our Lumi4-Tb donor together with green and red acceptors (**Figure S3**). pAz was previously reported to increase labeling efficiency through stabilization of a pre-complex with copper^26^. Secondly, we used 2-[4-{(bis[(1-tert-butyl-1H-1,2,3-triazol-4-yl)methyl]amino)methyl}- 1H-1,2,3-triazol-1-yl] acetic acid (BTTAA) as a ligand to stabilize Cu(I) in solution^27^. Indeed, we found that BTTAA outperformed tris[(1-hydroxy- propyl-1H-1,2,3-triazol-4-yl)methyl]amine (THPTA) with regard to reaction efficiency (**Figure S2b**). We further tested different copper concentrations with regard to cytotoxicity and labeling efficiency. No significant cell death was found at 200-500 µM CuSO4 in the presence of 6 equivalents of BTTAA (**Figure S2c**) and labeling was proceeding with similar efficiency under these conditions (**Figure S2d**). We further found that ratios of 5-6 equivalents BTTAA over CuSO4 provided better labeling specificity compared to lower ratios (**Figure S2e**). We then tested the reactivity of the incorporated PrF towards our pAz dyes at increasing concentrations and found that reaction with pAz-Lumi-4Tb (**Figure S4a**) was proceeding slower than for the pAz-green (**Figure S4b**) and pAz-red (**Figure S4c**) acceptors. Nevertheless, all reactions nearly proceeded to completion as judged by comparison of receptors labeled at N-terminal SNAP-tags (**Figure S4a-c**). More importantly, consecutive labeling of the SNAP-tags with Lumi4-Tb donor followed by CuAAC to combine it with either green or red acceptors resulted in clean FRET signals between the upper and lower VFT lobes, significantly exceeding background signals and reaching saturation at only a few µM of pAzF dye (**Figure S4d-f**).

Using these optimized conditions, we performed live HEK293T cell LRET imaging of mGlu2, labeled with SNAP-Lumi4-Tb via the SNAP-tags and PrF248 with pAz-green via CuAAC (**Figure 1d**). This confirmed the live cell-compatibility of our final labeling protocol, with a significant FRET signal as a result of donor-sensitized acceptor emission between the upper and lower lobes of the VFTs.

### LRET-based conformational sensors of the extracellular domain for investigating mGlu2 receptor activation

We further verified the functionality of SNAP-mGlu2-PrF248 by IP (phosphoinositol monophosphate) accumulation in response to different orthosteric agonists. It showed native-like pharmacological profiles (**Figure S5a-b**) and nearly identical EC50 values for a set of different ligands as compared to the wildtype receptor (**Figure 1e**). Titration of the corresponding doubly-labeled receptors with the same set of ligands revealed dose-dependent increases in LRET as a response to the expected closure of the VFTs bringing the upper lobe and the lower lobe into closer proximity (**Figure 1f**). These titrations also demonstrated that agonist efficacy and potency can be recovered by monitoring VFT closure directly at the level of receptor conformation, with results similar to IP-One accumulation, reporting on a downstream cellular response (**Figure 1g**).

Altogether, we established a procedure for the incorporation and efficient labeling of a ncAA in mGlu2, while maintaining cell viability, allowing to perform LRET measurement in live cells together with functional assays. This protocol provides easy access to screen other labeling positions, in order to design, test, and validate new conformational sensors for mGlu2 that can subsequently be solubilized to perform smFRET experiments.

### Monitoring mGlu2 VFT closure in a single protomer using dual ncAA labeling

The combination of N-terminal SNAP-tag labeling and incorporation of a single PrF residue in the lower lobes of each VFT within the dimeric receptor allowed us to monitor VFT closure by LRET in response to different orthosteric agonists. This encouraged us to circumvent the use of the ∼19.4 kDa large SNAP-tag for labeling by incorporating a second PrF residue within the upper lobe of the VFT. For this, we tested multiple sites within the VFTs upper lobe in combination with 248-TAG for double PrF incorporation efficiency and functional integrity of receptors by IP-One accumulation (**Figure S1**). Based on this screening we found a mutant with PrF at positions 358 in the upper and 248 in the lower lobe of each of the VFT protomers (SNAP-mGlu2- PrF248+358), outcompeting other pairs with regard to yields (**Figure S1**). Its pharmacological profile resembled the one obtained for wildtype receptors (**Figure S5a and c**), with a good correlation of EC50 values for different ligands (**Figure 2a**). Likewise, LRET measurements of this mGlu2-PrF248+358 sensor, stochastically labeled with pAz-Lumi4Tb donor and pAz-green acceptor, reported on the ligand-induced closure of the VFTs in an intrasubunit fashion (**Figure 2b**) and EC50 values correlated well with IP accumulation measurement (**Figure 2c**).

**Figure 2:**
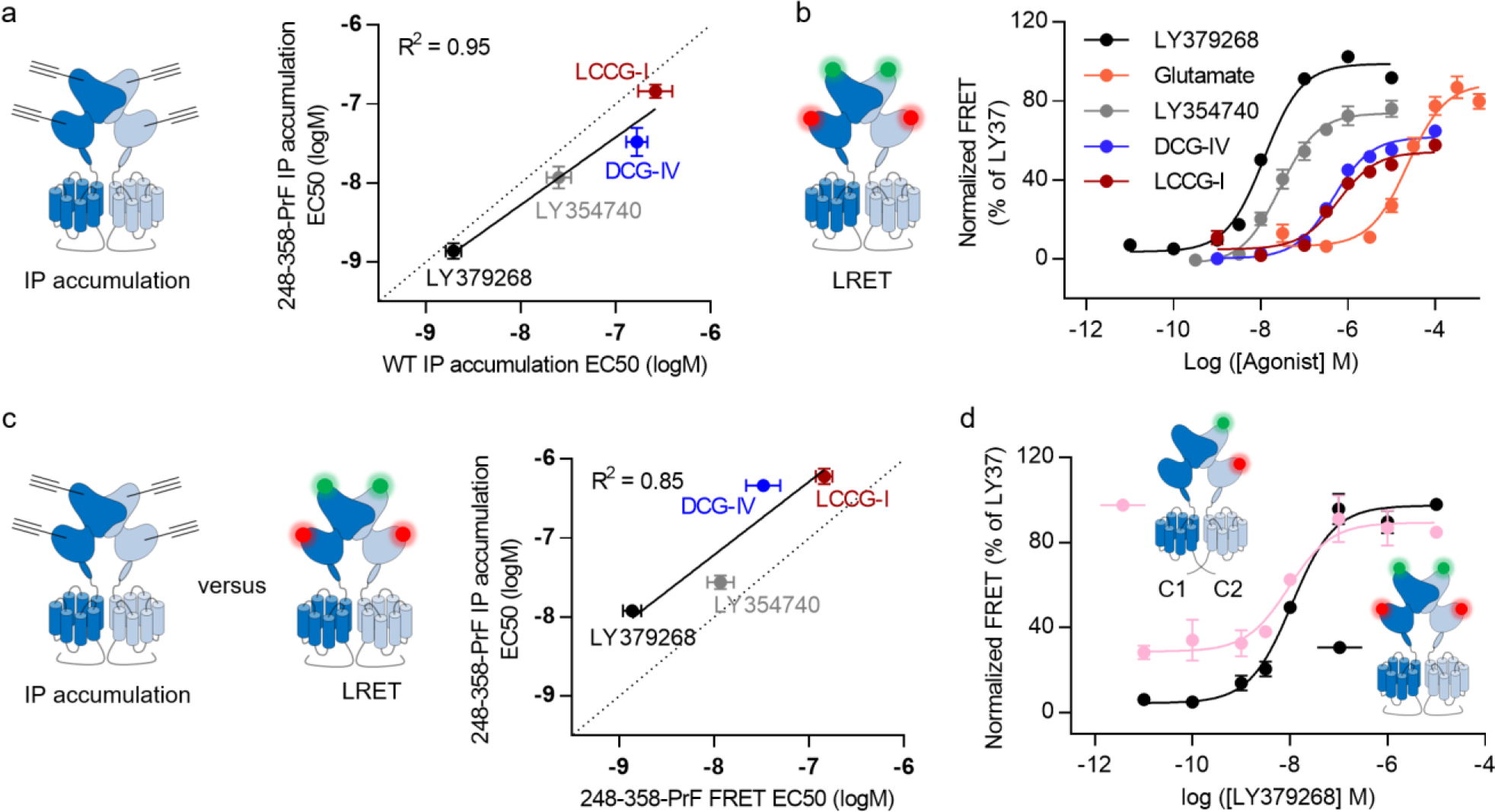
Establishment of a sensor to monitor VFT closure in a single protomer. a) Comparison of logEC50 values for different ligands obtained on WT mGlu2 and mutant with incorporated PrF at R358 and A248 on the upper and lower lobes of the VFTs in both subunits, respectively. b) Dose-response curves reporting on VFT closure in response to different ligands monitored by LRET. Both subunits were stochastically labeled with Lumi4Tb-pAz donor and green-pAz acceptor by click chemistry with incorporated PrF at R358 and A248. c) Comparison of logEC50 values for different ligands, determined by IP accumulation and LRET. d) Comparison of LRET changes upon titration with the full agonist LY379268 in constructs containing the VFT FRET sensor in one (purple) or both receptor subunits (black). Labeling of only one subunit was achieved using the previously described GABAB quality control system, which allows only heterodimers of PrF248-PrF358-C1 and WT-C2 C-terminal tails to be transported to the cell surface. Data in e-g represent the mean of 3-12 measurements +/- SD.

Finally, in order to obtain individual mGlu2 dimers carrying a single donor and a single acceptor fluorophore, as needed for smFRET measurements, we generated receptors with ncAAs incorporated in the upper and lower lobes of only one protomer. We took advantage of the engineered GABAB quality control system we have previously developed^28^ to specifically label heterodimers at the cell surface, containing PrF only in the upper and lower lobes of one VFT protomer (SNAP-mGlu2-PrF248+358-C1/SNAP-mGlu2-C2). We compared the changes in LRET induced by the full agonist LY379268 on VFT closure in both variants, and found strikingly similar results with regard to LY379168 potency, while only slight differences in the low and high average LRET efficiencies were observed (**Figure 2d**). Such heterodimers allowing to monitor the VFT closure from a single FRET pair within one protomer are suitable for our confocal single molecule experiments and further rule out contributions arising from intersubunit FRET.

### Full agonists induce maximal VFT closure independent of allosteric modulation

While LRET is powerful to determine ligand potency and efficacy in a population of receptors, identifying individual conformational states requires single molecule analysis. Therefore, we installed the smFRET- compatible Alexa488 (donor) and Alexa647 (acceptor) in a stochastic manner on the SNAP-mGlu2-PrF248+358- C1/SNAP-mGlu2-C2 sensor (**Figure 3a**, in the following denoted as U358-L248 sensor). As this strategy requires the modification of the C-terminal tails of mGlu2 we first verified the functionality of these constructs after solubilization. To account for the decreased yields of receptors with PrF incorporated by genetic code expansion, we modified our previously published solubilization protocol^17^. This new strategy relies on an initial solubilization using 1% LMNG + 0.1% CHS Tris followed by dilution into buffer containing GDN (as compared to our previously described protocol using 0.1% LMNG + 0.01% CHS Tris + 0.1% GDN for initial solubilization followed by dilution in buffer without additional detergent). Control LRET titration experiments on SNAP-tag labeled receptors confirmed unimpaired functional integrity over time at room temperature and demonstrated that the C-terminal modifications do not impair receptor stimulation by Glu and allosteric modulation by BINA (**Figure S6a-d**). In addition, the same constructs were analyzed by smFRET, which confirmed that intersubunit VFT reorientation occurs in an identical manner on these C-terminally modified constructs as on wildtype receptors (as in ^17^) in the absence of ligands or when saturated with LY341495, Glu, Glu + Gi and Glu + BINA (**Figure S7a-e**).

**Figure 3:**
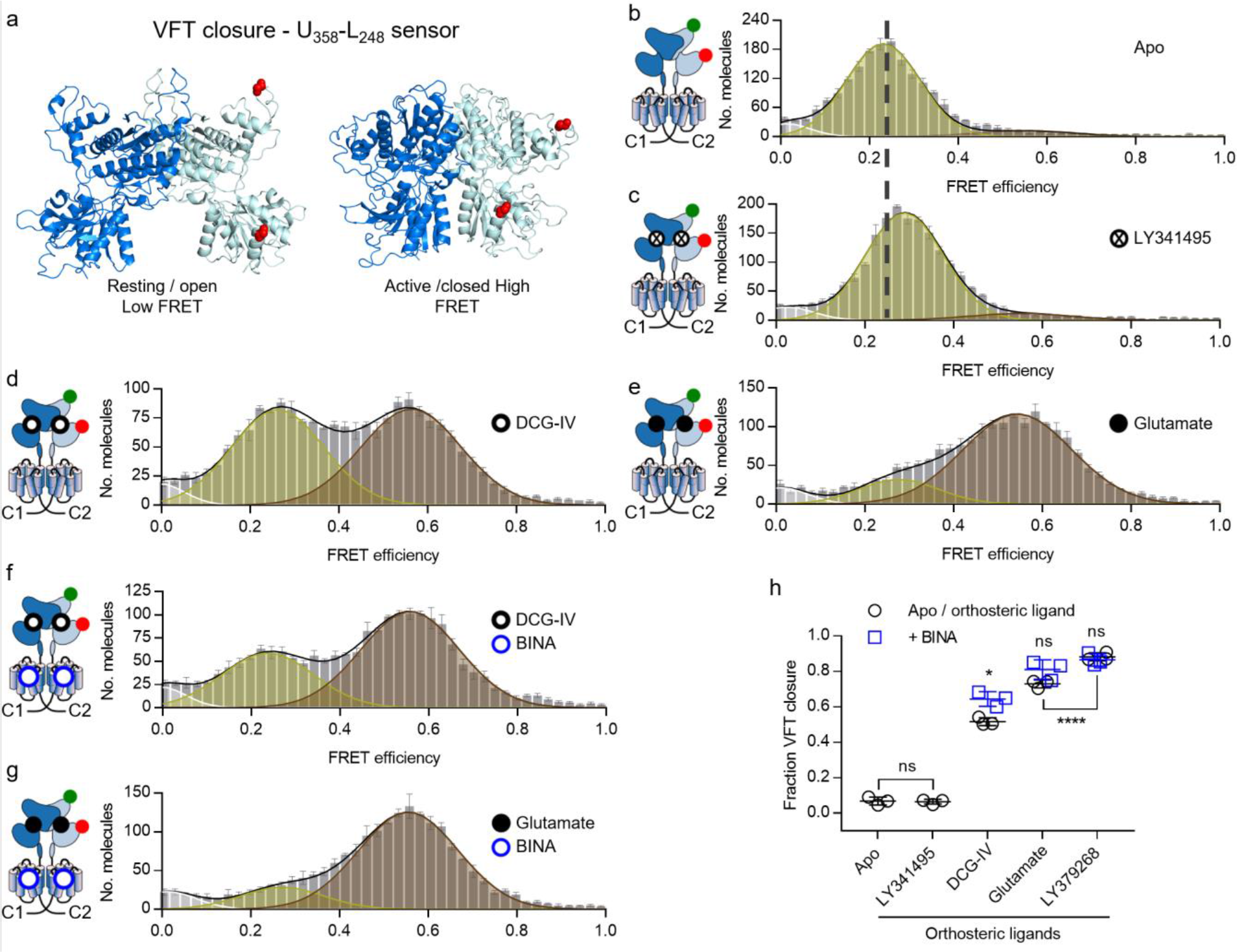
Agonists strongly increase the residence time of the VFT in its closed state. a) Side-view on structures of the VFT in the inactive resting open/open (PDB ID 7EPA) and active closed/closed (PDB ID 7E9G) conformation with residues R358 in the upper lobe and A248 in the lower lobe of one VFT replaced by PrF highlighted in red spheres. b-g) FRET histograms of VFT closure sensor (UL) in the absence of ligand (b, Apo) and in the presence of saturating LY341495 (c), DCG-IV (d), DCG-IV + 1 µM BINA (e), Glutamate (f) and Glutamate + 1 µM BINA (g). FRET histograms in b-g show the accurate FRET efficiency as the mean +/- SEM of three independent biological replicates each normalized to 2000 events in the DA population (S = 0.3-0.7). Histograms display the fitting with 3 gaussians (white = very low FRET, yellow = low FRET, red = high FRET) together with the global fit (black). The mean FRET efficiency of the Apo condition (a) in comparison to the antagonist LY341495 (b) is indicated by a dashed line. h) Scatter plot showing the fraction of molecules in the closed state. Shown are the means +/- SD of three independent biological replicates obtained from the number of molecules found in the high FRET population (red) over the sum of all molecules found in the low FRET (yellow) and high FRET (red) populations. Black circles show the Apo condition and orthosteric ligands alone while blue squares are given for agonists in the presence of 1µM BINA. Statistical differences were determined using a one-way ANOVA with Sidak multiple comparisons test and are given as ****p ≤ 0.0001, ***p ≤ 0.001, **p ≤ 0.01, *p ≤ 0.05, ns > 0.05.

The FRET distribution of our U358-L248 sensor showed a major population at a low FRET value of 0.23 in the absence of ligand, indicating that the open conformation is strongly favored under this condition **(Figure 3b)**. Addition of a saturating concentration of the antagonist LY341495 led to a similar profile with a slight shift of the population toward a higher FRET value of 0.29 **(Figure 3c)**. In contrast, partial agonist DCG-IV led to the appearance of a second population, characterized by a high FRET state around 0.56 **(Figure 3d**), whose relative population was further increased by the PAM BINA **(Figure 3f)**. Glutamate led to a strong population of the high FRET state **(Figure 3e)** with a nearly complete depopulation of the low FRET state, while BINA had no additional effect **(Figure 3g)**. Similar results were obtained for LY379268 **(Figure S8a-b)**. Note that we excluded the possibility of significant contaminations in the observed FRET populations with undesired dimers composed of receptors being labeled on both VFTs **(Figure S8c-d)**. We excluded the minor population at very low FRET (E = 0.01) **(Figure 3b-g and S8a-b)** from further analysis, since it was not sensitive to ligands **(Figure S8e)**.

In accordance with structural data (**Figure 3a**), we assigned the agonist-induced high FRET population to the closed conformation and the low FRET population to the open conformation of the VFT **(Figure 3b-g**). We then quantified the fraction of VFT closure, i.e. the number of molecules found in the closed population over the sum of all molecules found in both populations **(Figure 3h)**. Statistical analysis confirmed no difference in VFT closure between apo and the antagonist LY341495. For other ligands, the fraction of closed VFTs reflected agonist efficacy, following DCG-IV < Glutamate < LY379268, while the PAM only had a significant effect on the partial agonist DCG-IV and not on the two full agonists.

The closed state of the VFT can be described by a single distance, independent of which ligand is bound with a mean FRET efficiency centered around E = 0.57. On the contrary, differences for the low FRET state were observed for the different conditions (**Figure 3b-g and S8a-b**). This indicates that ligands led to slightly different conformations of the open state, in a way that seemed uncorrelated to their relative efficiency in promoting VFT closure or their pharmacological profile (**Figure S8f**). Overall, the coexistence of a low FRET and a high FRET population proposes a dynamic equilibrium between open and closed conformations, where agonist action stems from their ability to stabilize the closed conformation.

### Reorientation of the VFT upper lobes is favored by allosteric modulation

To monitor the reorientation of the mGlu2 VFT dimer at the level of the upper (U) lobes, we incorporated PrF at Arg358 and labeled it with Alexa546 (donor) and Alexa647 (acceptor) (U358-U358 sensor) **(Figure 4a)**.

**Figure 4:**
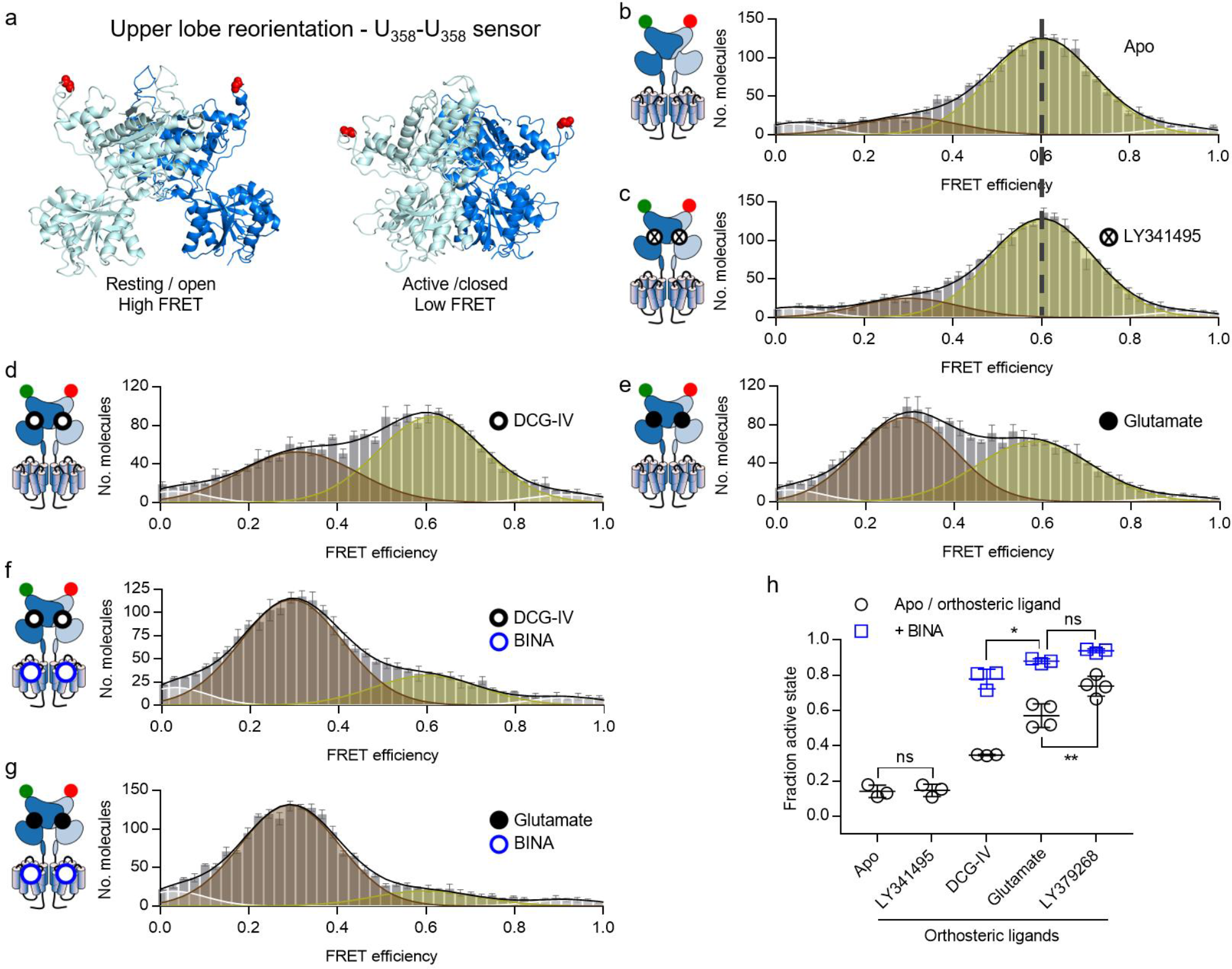
The equilibrium between resting and active upper lobe conformation follows agonist efficacy and is further promoted by allosteric modulations. a) Side-view on structures of the VFT in the inactive resting closed/closed (PDB ID 7EPA) and active open/open (PDB ID 7E9G) conformation with residues R358 replaced by PrF highlighted in red spheres. b-g) FRET histograms of upper lobe sensor in the absence of ligand (b, Apo) and in the presence of saturating LY341495 (c), DCG-IV (d), Glutamate (e), DCG-IV + BINA (f), and Glutamate + BINA (g). FRET histograms in b-g show the accurate FRET efficiency as the mean +/- SEM of three independent biological replicates each normalized to 2000 events in the DA population (S = 0.3-0.7). Histograms display the fitting with 4 gaussians (white = very low FRET, yellow = low FRET, red = high FRET, white = very high FRET) together with the global fit (black). The mean FRET efficiency of the Apo condition (a) in comparison to the antagonist LY341495 (b) is indicated by a dashed line. h) Scatter plot showing the fraction of molecules in the fully reoriented state. Shown are the means +/- SD of three independent biological replicates obtained from the number of molecules found in the high FRET population (red) over the sum of all molecules found in the low FRET (yellow) and high FRET (red) populations. Black circles show the Apo condition and orthosteric ligands alone while blue squares are given for agonists in the presence of 10µM BINA. . Statistical differences were determined using a one-way ANOVA with Sidak multiple comparisons test and are given as ****p ≤ 0.0001, ***p ≤ 0.001, **p ≤ 0.01, *p ≤ 0.05, ns > 0.05.

FRET histograms showed a multimodal distribution that was best accounted for by fitting with 4 gaussians (**Figure 4b-g and S9a-b**). A major high FRET population was found at E = 0.6 in the absence of ligand and presence of LY341495 **(Figure 4b-c).** It was partially depopulated in the presence of DCG-IV, resulting in the appearance of a second major low FRET population centered around E = 0.3 (**Figure 4d**). Furthermore, BINA promoted an increase in the population of the low FRET state **(Figure 4e)**. Glutamate had a similar but more efficient effect than DCG-IV on VFT upper lobe reorientation both in the absence and the presence of the BINA **(Figure 4f-g)**, while LY379268 was even more efficient **(Figure S9a-b)**. To quantify the reorientation of the VFTs upper lobes, we first determined the mean FRET efficiency E and the width of the FRET distribution (FWHM) values for the four populations under different ligand conditions. We found only minor deviations of recovered mean E and FWHW between the different conditions and therefore fixed these values to quantify the population of the four states. As we did not observe ligand-related changes in the population of the very low and very high FRET states (**Figure S9c-d**, E = 0.03 and 0.92, respectively), we annotated the fraction of upper lobe VFT reorientation as the number of molecules in the low FRET population over the sum of molecules found in the low and high FRET populations **(Figure 4h)**. The results confirmed a ligand-dependent impact on the equilibrium of VFT reorientation with Apo = LY341495 < DCG-IV < Glu < LY379268. In addition, BINA revealed differences in the maximal reorientation according to ligand efficacy

Thus, upper lobe VFT reorientation can be described by an equilibrium between the inactive high FRET and the active low FRET states, where orthosteric agonists exert differences in efficacy by shifting this equilibrium towards the active conformation, without stabilizing distinct states. In contrast, the PAM that binds in the 7TM domain is strictly required to stabilize the fully reoriented VFTs in the active state, even though this effect to increase efficacy remains dependent of the orthosteric agonist. We note that these observations are perfectly consistent with those we previously obtained using SNAP-tagged receptors, labeled at the level of the upper lobes as well ^17^.

### The lower lobe VFT reorientation limits ligand efficacy but is favored by an allosteric modulator

Next, we created a VFT intersubunit sensor reporting of the reorientation of the lower (L) lobes by installing the same donor and acceptor fluorophores after incorporation of PrF at Ala248 (L248-L248 sensor) **(Figure 5a)**.

**Figure 5:**
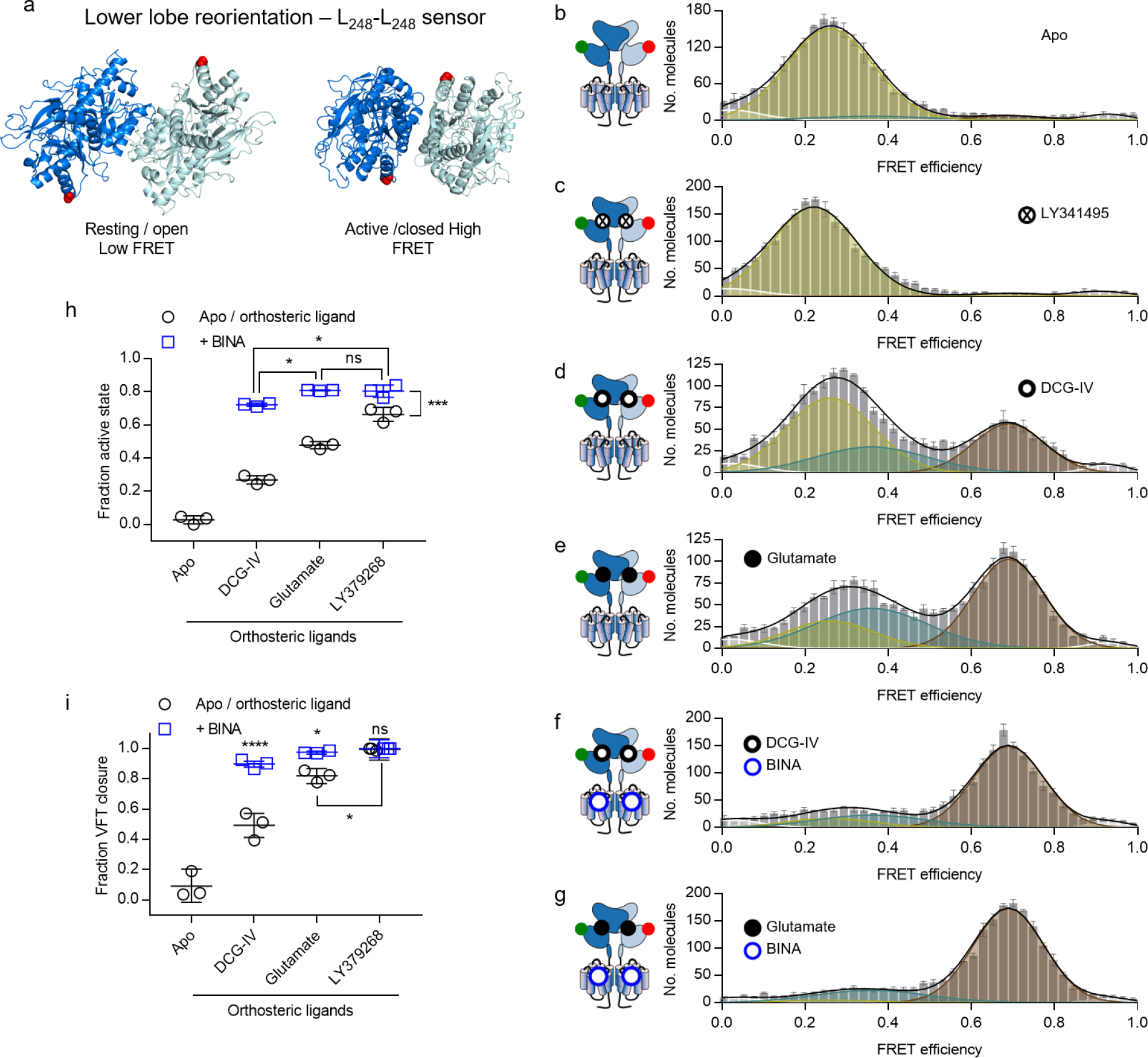
The lower lobes of the VFT report on VFT closure and activation. a) Bottom-view on structures of the VFT in the inactive resting open/open (PDB ID 7EPA) and active closed/closed (PDB ID 7E9G) conformation with residues A248 replaced by PrF highlighted in red spheres. b-g) FRET histograms of lower lobe sensor in the absence of ligand (Apo, b) and in the presence of saturating LY341495 (c), DCG-IV (d), Glutamate (e), DCG-IV + BINA (f) and Glutamate + BINA (g). FRET histograms show the accurate FRET efficiency as the mean +/- SEM of three independent biological replicates each normalized to 2000 events in the DA population (S = 0.3-0.7). Histograms display the fitting with 5 gaussians (white = very low FRET, yellow = low FRET1, blue = low FRET2, red = high FRET, white = very high FRET) together with the global fit (black). h) Scatter plot showing the fraction of molecules in the fully reoriented/active state. Shown are the means +/- SD of three independent biological replicates obtained from the number of molecules found in the high FRET population (red) over the sum of all molecules found in the low FRET1 (yellow), low FRET2 (blue) and high FRET (red) populations. i) yellow = low FRET1, blue = low FRET2, red = high FRET, white = very high FRET) together with the global fit (black). h) Scatter plot showing the fraction of molecules in the closed VFT state. Shown are the means +/- SD of three independent biological replicates obtained from the sum of molecules found in the high FRET (red) and low FRET2 (blue) populations over the sum of all molecules found in the low FRET1 (yellow), low FRET2 (blue) and high FRET (red) populations. h-i) Black circles show the Apo condition and orthosteric ligands alone while blue squares are given for agonists in the presence of 10µM BINA. Statistical differences were determined using a one-way ANOVA with Sidak multiple comparisons test and are given as ****p ≤ 0.0001, ***p ≤ 0.001, **p ≤ 0.01, *p ≤ 0.05, ns > 0.05.

From the recent cryo-EM structures of mGlu2 in the inactive and active states(Du et al., 2021; Lin et al., 2021a; Seven et al., 2021), we predicted a shift from a low FRET to a high FRET signal upon receptor activation.

Accordingly, a major population centered around E = 0.25 was obtained for the apo receptor **(Figure 5b)**. A saturating concentration of LY341495 slightly but significantly reduced this mean E to 0.22, indicating a further separation of the lower lobes **(Figure 5c)**. Agonists led to the appearance of a high FRET population centered around E = 0.69, a value that remained constant for different agonist conditions, pointing at a single distance between the lower lobes in the active, reoriented state (**Figures 5d-g and S10a-h)**. In all cases, the low FRET (LF) population remained significantly populated, but its FRET value, reporting on the distance between the lower lobes, varied significantly between 0.25 to 0.35 (**Figures 5d-g and S10a-h).** In contrast to what was observed with the U358-U358 sensor, a correlation between the efficacy of the ligands and the mean FRET efficiency of the Low FRET (LF) state was found for the L248-L248 sensor **(Figure S10i)**. We propose that this shift in LF value could stem from the transition of the receptor from a Resting-open (Ro) to a Resting-closed (Rc) state, for which the distance between A248 would differ from only a few angstroms, according to VFT crystal structures obtained on mGlu3 ^10^. In the presence of antagonist, the Ro conformation is preferentially populated (With ELF = 0.25), while with LY379268 the receptor is mainly in the closed conformation (with ELF= 0.35). Accordingly, we fitted our histograms with 3 main populations assigned to the Ro, Rc and A conformations (with E = 0.25 (yellow), 0.35 (blue) and 0.69 (red) respectively, **Figures 5b-g** and **S10j-k**). Then we calculated the fraction of receptors in the active state (**Figure 5h**) and in the closed state (**Figure 5i**) (as A/(Ro+Rc+A) and (Rc+A)/(Ro+Rc+A), respectively). For the different ligands, these calculated values with this L248-L248 sensor are remarkably similar to the ones observed on the U358-L248 and U358-U358 sensors, that report mainly on VFT closure and VFT dimer reorientation, respectively (Compare **Figures 3h and 5i**, and **4h and 5h**). This observation therefore validated our interpretation of the complex smFRET data obtained with this set of complementary sensors.

## Discussion

Here we present a live cell compatible click chemistry approach to site-specifically label mGlu receptor protomers with donor and acceptor fluorophores. We demonstrate that it allows to screen for suitable positions to establish FRET sensors, directly reporting on conformational rearrangements of individual receptor domains by LRET and smFRET. Although a set of different ncAAs together with corresponding bioorthogonal chemistries are nowadays available to achieve a site-directed fluorescence labeling of proteins, the CuAAC remains one of the fastest and most specific reactions^29^. Nevertheless, the need of copper has clearly limited its use for labeling of proteins directly within the membrane of living cells^30^. Through careful optimization and the combination of the advanced copper-chelating ligand BTTAA with pAz dyes we achieve an efficient labeling of cell-surface receptors within only 25 min at 37°C, while maintaining cell viability. This approach opens up new possibilities to study ligand pharmacology, by directly looking at their effect on individual domain rearrangements. Combined with classical pharmacological assays such as the accumulation of IP, this allows to link GPCR conformational changes to downstream cellular responses. While our ensemble LRET approach allows to efficiently screen for the effect of different ligands, we further show that suitable FRET sensors can subsequently be used in smFRET measurements after receptor solubilization to dissect the underlying conformational states and dynamics.

Here, we focused on the initial step of mGlu2 activation, e.g. the major inter- and intrasubunit rearrangements occurring within the VFTs of this dimeric class C GPCR. Through the analysis of these conformational rearrangements on single receptors solubilized in proper detergents, we are able to show that mGlu2 VFT activation is governed by conformational equilibria between different states. Overall our data point to a model, where orthosteric and allosteric ligands act differently on an equilibrium between three states, corresponding to the Ro, Rc and A conformations (**Figure 6**). Orthosteric ligands appear to act on the open/closed equilibrium (Ro / Rc). They promote and stabilize the closure of the VFT domain to an extent that is correlated with their pharmacological efficacy. The three tested agonists (DCG-IV, glutamate and LY379268) lead to a high degree of VFT closure (52%, 73% and 88% respectively, **Figure 3h**), while their ability to reorient the receptor to the active state is more limited (35%, 57% and 74% respectively, **Figure 4h**). The PAM BINA acts on a second equilibrium, between the Rc and A conformations. In its presence, the Rc state is destabilized, the receptor reorientation toward its active state is favored, and therefore the closure of the VFT by a ligand is more directly linked to the reorientation. Accordingly, the correlation between the extent of VFT closure (measured by the U358-L248 sensor (**Figure 3h**)) and the dimer reorientation ((measured by the U358- U358 sensor (**Figure 4h**) increases in the presence of the PAM (for the 3 agonists, 64%, 81%, 86% of VFT closure vs. 78%, 88%, 94% reorientation).

**Figure 6:**
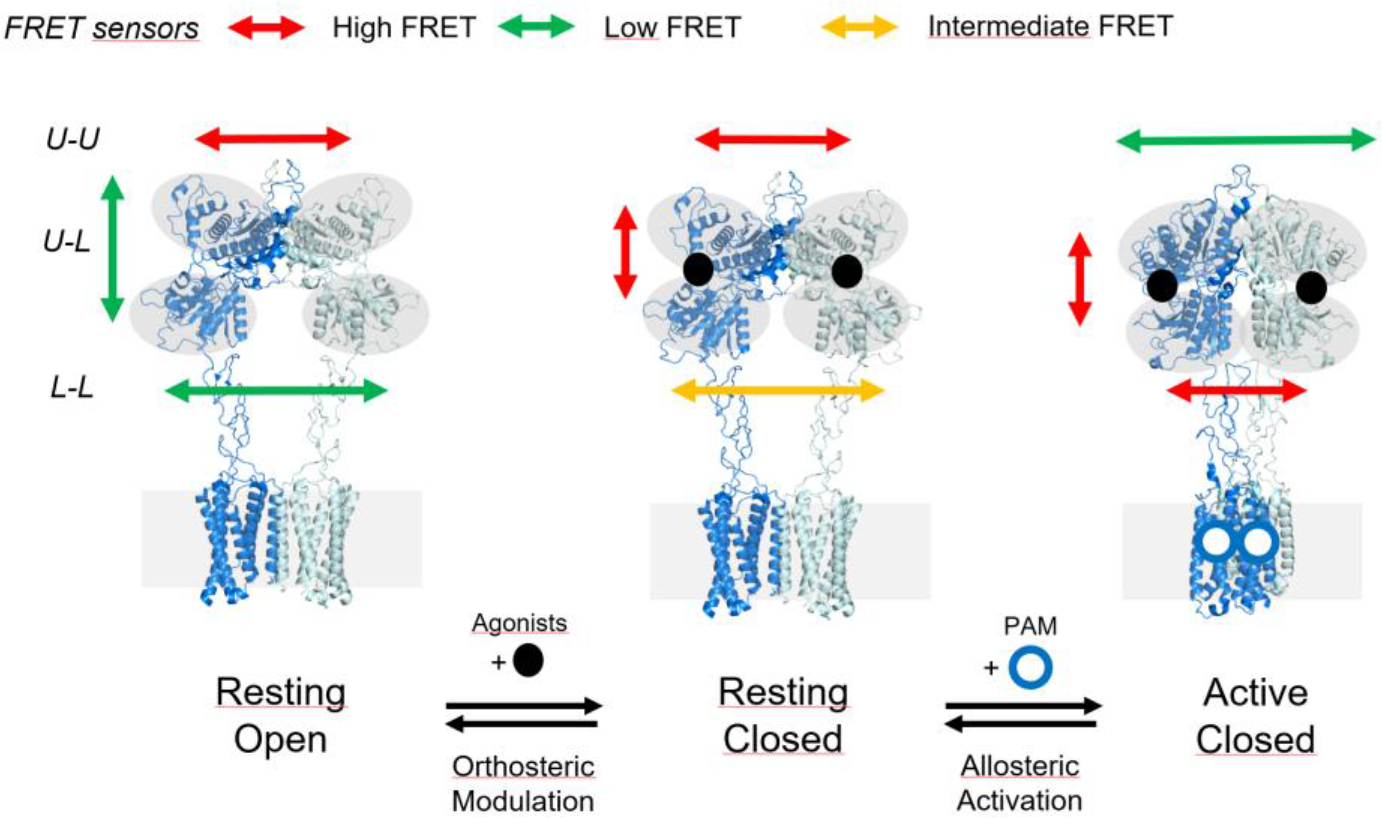
3-state model of mGlu2 activation. The initial step in mGlu2 activation can be described by a 3-state equilibrium between the Resting Open, the Resting Closed and the Active Closed states. Orthosteric ligands act on the Ro-Rc equilibrium, where agonists populated the Rc state in a manner reflecting their pharmacological efficacies. In the absence of allosteric modulators, the VFTs samples between the Rc and Ac states. Positive allosteric modulators act on the Rc-A equilibrium by increasing the residence time of molecules in the Ac state.

Interestingly, the variations in the mean FRET efficiency of our VFT closure sensor (**Figure 3**) indicate the adoption of distinct open conformations for the different orthosteric ligands tested, but not being correlated with agonist efficacies. In contrast, the closed state is represented by a single average donor/acceptor distance and thus likely a highly similar single conformation no matter what agonist is bound. This points at ligand-induced conformational changes and further underlines the idea that the VFT constantly oscillates between distinct open and a single fully closed state when bound to ligands. We further identify an additional antagonist-bound state, which is characterized by a slight decrease in the distance between upper and lower lobes (**Figure 3c**) and an increase in the lower lobe separation (**Figure 5c**). These differences correlate well with the two recently described inactive mGlu2 structures, where the distance between the Cα atoms of our VFT closure sensor shows a 2.1Å decrease and the lower lobe distance increases by 7.1Å, when bound to antagonist^14^ as compared to the ligand-free VFT^15^.

Overall, our data demonstrate that the combination of genetic code expansion and click chemistry allows to establish FRET sensors to recover structural information with precision of a few Å and map the conformational landscape of GPCRs under various different conditions using smFRET. In addition, it provides access to the underlying conformational dynamics and how these are influenced by orthosteric and allosteric modulators. This enables to complement high resolution structural information obtained on stabilized receptors to dissect their mechanism of activation.

## Materials and Methods

### Materials

Aminoguanidine hydrochloride, Copper(II) sulfate (CuSO4), (+)-sodium L-ascorbate, Tris[(1-benzyl-1*H*- 1,2,3-triazol-4-yl)methyl]amine (TBTA), Tris(3-hydroxypropyltriazolylmethyl) amine (THPTA), L-glutamate and chemicals used for synthesis were purchased from Sigma-Aldrich (St. Louis, MO, USA) unless stated otherwise. LY379268, LY354740, DCG-IV, L-CCG-I, and BINA were purchased from Tocris Bioscience (Bristol, UK). O6-benzylguanine and amine derivatives of Lumi4-Tb, “green” (fluorescein derivative), ”red” (Cy5 derivative) fluorophores as well as Tag-lite® buffer were from Cisbio Bioassays (Codolet, France). 4- Propargyloxy-L-phenylalanine (PrF) was initially synthesized according to ^21^ and later purchased from Iris Biotech GmbH (Marktredwitz, Germany). 2-[4-{(bis[(1-tert-butyl-1H-1,2,3-triazol-4-yl)methyl]amino)methyl}- 1H-1,2,3-triazol-1-yl] acetic acid (BTTAA) was synthesized as described in ^27^. pAz-Alexa488, pAz-Alexa546 and pAz-Alexa647 were purchased from Jena Bioscience (Jena, Germany). Lauryl maltose neopentyl glycol (LMNG), and cholesteryl hemisuccinate (CHS) tris salt were purchased from Anatrace (through CliniSciences, France). Glyco-diosgenin (GDN) was purchased from Avanti Polar Lipids through Merck.

### Synthesis of Picoly-azide dye derivatives

Picolyl-azide (pAz) derivatives of Lumi4-Tb, “green” (fluoresceine) and “red” (Alexa647) fluorophores, were synthesized according to ^26^ with minor modifications as specified below. 2,5-Pyridinedicarboxylic acid was first esterified to 2,5-Pyridinedicarboxylic acid dimethyl ester in 10 eq. methanol by addition of 0.5 eq. concentrated H2SO4 through stirring for 18 h under reflux. After neutralization with a saturated aqueous solution of NaHCO3 and extraction with chloroform, the combined organic phases were dried over MgSO4, filtered, and the solvent removed under reduced pressure to yield the diester. 6-Azidomethylnicotinic acid was then prepared as described in ^26^ and subsequently converted to an N-Hydroxysuccinimid ester in Dimethylformamide through addition of 1.2 eq. N,N,N′,N′-Tetramethyl-O-(N-succinimidyl)uronium tetrafluoroborate followed by 6 eq. Diisopropylethylamine. After 1 h of stirring, formation of the product was verified by LC/MS, the solution was filtered (0,2 µm PTFE membrane, GE Healthcare, France) and directly added to fluorophore amines at 1.7 eq. in DMSO. 6 eq. of Diisopropylethylamine were added and the solution stirred for 2 h. pAz dye conjugates were subsequently purified by HPLC and product formation was confirmed by LC/MS (pAz-Lumi4Tb, 60%, C63H74N17O13Tb: theoretic mass = 1435.4906 Da, experimental mass = 1435.4906 Da; pAz-green, 64%, C36H33N7O8: theoretic mass = 691.1391 Da, experimental mass = 691.2391 Da; pAz-red, 1.6%, C41H49N8O8S2: theoretic mass = 845.3109 Da, experimental mass = 845.3109 Da).

### Plasmids

The pcDNA plasmid encoding human mGlu2 with N-terminal FLAG- and SNAP-tags was a gift from Cisbio Bioassays (Perking Elmer, Codolet, France). The pRK5 plasmid encoding the rat SNAP-mGlu2 and the construction of the rat SNAP-mGluR2-C1KKXX and SNAP-mGluR2-C2KKXX plasmids were described in^31^. Premature Amber mutations were introduced using the QuickChange Multi Site-Directed Mutagenesis Kit from Agilent Technologies (Santa Clara, CA, USA) according to the manufacturers protocol. The bicistronic pcDNA vector coding for the engineered *E. coli* tyrosly-tRNA synthetase (PrFRS, pPR-EcRS2 according to ^21^ and including the enhancing mutation R265 according to ^22^) and three copies of the engineered *B. stearothermophilus* tyrosyl-tRNACUA^23^ was generated starting from the corresponding pET21 plasmid provided by Pierre Paoletti’s lab (IBENS, ENS, Paris, France). The generation of the pRK5 plasmid coding for the high-affinity glutamate transporter EAACI was described in^32^. Samples for smFRET experiments were prepared using the pIRE4-PGK- EPrFRS kindly provided by Irene Coin (obtained by mutagenesis of the aminoacyl-tRNA synthetase to Thr37, Ala183, Leu186 in the pIRE4-Azi plasmid described in ^33^).

### Cell culture, transfection and incorporation of UAA

Adherent HEK293T cells (ATCC CRL-3216, LGC Standards S.a.r.l., France) were cultured in DMEM (Thermo Fisher Scientific) supplemented with 10% fetal bovine serum (FBS, Sigma-Aldrich) at 37°C with 5% CO2. HEK293T cells were transfected using Lipofectamine® LTX with Plus™ Reagent (Thermo Fisher Scientific) in 96-well F-bottom black plates (Greiner Bio-One). 45,000 cells were seeded in 100 µL DMEM per well 24h before transfection. Transfections were performed using 255 ng total DNA per well with a 10:6:1 ratio of vectors coding for mGluR2:PrFRS-tRNACUA:EAACI in 30 µL Opti-MEM® I reduced serum medium (Thermo Fisher Scientific). 0.255 µL Plus™ Reagent per well were added and incubated for 10 min at RT in the dark. Then 0.75 µL Lipofectamine® LTX per well were added, followed by an additional incubation for 30 min at RT in the dark. Finally, the DNA-lipid complexes were added to the cells and incubated at 37°C for 6 h. Then the medium was replaced with 100 µL fresh DMEM supplemented with 0.5 mM PrF. The medium was further exchanged with fresh PrF-containing medium after 24 h and 48 h.

For tr-FRET microscopy, 100,000 cells per well were seeded in Nunc™ Lab-Tek™ chambered coverglass 8 well plates (Thermo Fisher Scientific) 24 h prior to transfection. Transfection and expression were carried out as described above but using 80 µL of total transfection mixture, which were added to 200 µl DMEM per well. 72 h post-transfection, medium was exchanged to DMEM supplemented with GlutaMAX™ (Thermo Fisher Scientific) without PrF and incubated for 2 h at 37°C and 5% CO2 before labeling.

For single molecule FRET measurements, 500,000 cells per well were seeded in 2 ml Gibco™ DMEM, high glucose, GlutaMAX™ Supplement, pyruvate (Thermo Fisher Scientific, France) + 10% (vol/vol) FBS (FBS, Sigma-Aldrich, France) on 6 well plates 24 h prior to transfection. Transfections were carried out using a total of 2 µg per well at a 1:1 ratio of pRK5-rat-mGlu2:pIRE4-PGK-EPrFRS DNA, diluted in 200 µl JetPrime Buffer together with 4 µl of JetPrime transfection reagent (Polyplus-transfection SA, Illkirch-Graffenstaden, France). The transfection mixture was incubated at RT for 15 min before addition to the cells. After 6 h and 24 h the medium was exchanged to fresh DMEM GlutaMAX™ + 10% FBS containing 0.5 mM of PrF and expression continued until 48 h post-transfection. Expression of C-terminally modified rat mGluR2-248TAG-358TAG- C1KKXX and mGluR2-C2KKXX constructs together with pIRE4-PGK-EPrFRS was performed for 72 h with medium exchange after 6, 24 and 48 h at a 4:1:5 DNA ratio, respectively. 2 h prior to labeling, medium was exchanged to Gibco™ DMEM, high glucose without phenol red, supplemented with GlutaMAX™ and pyruvate (Thermo Fisher Scientific, France) and incubated at 37°C and 5 % CO2. Expression of receptors labeled via the N-terminal SNAP-tags for functional characterization after solubilization and smFRET measurements, was carried out as described above but using 4 µl of Lipofectamine 2000 (Thermo Fisher Scientific, France) per 2 µg of total pRK5-rat-SNAP-mGlu2 in a total of 200 µl OptiMEM medium for transfection. Expression was achieved in complete medium without PrF for a total of 48 h.

### IP measurements and cytotoxicity assay

IP accumulation was measured using the IP-One HTRF® assay kit (Cisbio Bioassays, Condolet, France), following the official protocol from the manufacturer. Cytotoxicity was determined by staining with 2 µg/ml propidium iodide together with 5 µg/ml Hoechst 33342 for 15 min followed a single wash with PBS. Detection of the fluorescence was done using an Infinite F500 spectrofluorometer (Tecan).

### SNAP-tag labeling for quantification and LRET measurements

Labeling with a single fluorophore was performed on adherent cells at 37°C for 1 h using the indicated concentrations of SNAP-Lumi4-Tb, SNAP-green or SNAP-red diluted in Tag-lite® buffer. Double labeling was performed using 100 nM SNAP-Lumi4-Tb and 60 nM SNAP-green. After labeling, cells were washed three times with Tag-lite® buffer.

### Labeling of PrF-containing receptors with pAz dyes for LRET

The final CuAAC labeling mixture was composed of 1.5 mM aminoguanidine hydrochloride, 2 mM BTTAA, 0.36 mM CuSO4 together with 2 mM (+)-sodium L-ascorbate in Tag-lite® buffer. Single labeling was achieved by subjecting cells to the labeling mixture containing either 3 µM pAz Lumi4-Tb, pAz-green, pAz-red. For double labeling 3 µM of Lumi4-Tb donor and 8-10 µM of either pAz-green or pAz-red acceptor were used. Labeling was carried out for 25 min at 37°C in the dark. Cells were subsequently washed three times with Tag-lite® buffer before addition of ligands. For double labeling through the SNAP-tag and PrF, cells were first labeled via the SNAP-tag using 300 nM SNAP-Lumi4-Tb or 100 nM SNAP-green, wash once with Tag-lite® buffer and then label via CuAAC as described above, followed by three washes before adding ligands.

### Ensemble fluorescence and LRET measurements

An Infinite F500 spectrofluorometer (Tecan) was used to measure the emission of Lumi4-Tb at 620 nm with a 50 µs delay after excitation at 340 nm using an integration time of 400 µs, geen at 520 nm after excitation at 485 nm using an integration time of 1000 µs and red at 665 nm after excitation at 610 nm using an integration time of 1000 µs. Sensitized acceptor emission from LRET between Lumi4-Tb and green was measured at 520 nm with a 50 µs delay after excitation at 340 nm using an integration time of 400 µs and between Lumi4-Tb and red at 665 nm with a 50 µs delay after excitation at 340 nm using an integration time of 400 µs^31^.

LRET measurements of PrF-containing sensors and the SNAP sensor after receptor solubilization were performed on a PHERAstar FS microplate reader (BMG LABTECH) as described^17, 34, 35^. After Lumi4-Tb donor excitation with a laser at 337 nm, the donor emission was collected at 620 nm and the sensitized acceptor emission at 520 nm for green or 665 nm for red in 5 µs intervals for a total of 2.5 s. LRET was calculated as the ratio of the sensitized acceptor emission integrated from 50 µs to 100 µs over 1200 µs to 1600 µs after donor excitation for the pure SNAP sensor and over 300 µs to 500 µs for SNAP with PrF and pure PrF sensors.

### LRET microscopy

The microscope used to image LRET was described previously^36^. In brief, “Green” was excited at 470-40 nm using a 495LP nm dichroic filter and imaged through a 520-15 nm filter, while “Lumi4-Tb” was excited at 350- 50 nm using a 400LP dichroic filter and imaged through a 550-32 nm filter. For LRET Lumi4-Tb was excited at 350-50 nm using a 400LP nm dichroic filter and the sensitized acceptor emission was collected using a 520-15 nm.

### Labeling, preparation of membrane fractions and receptor solubilization

Prior to solubilization from crude membranes, receptors were labeled for 1 h at 37°C with either 100 nM SNAP-Lumi4-Tb and 60 nM SNAP-green in Gibco™ DMEM, high glucose, GlutaMAX™ Supplement, pyruvate (Thermo Fisher Scientific, France) for LRET measurements or 600 nM BG-Cy3B and 300 nM SNAP- red in Gibco™ DMEM, high glucose without phenol red, supplemented with GlutaMAX™ and pyruvate (Thermo Fisher Scientific, France) for SNAP-sensor smFRET measurements. Labeling of incorporated PrF for smFRET measurements was achieved as described above using 2.5 µM pAz Alexa546 and 7.5 µM pAz Alexa647 (PrF358 upper lobe sensor and PrF248 lower lobe sensor) or 2.8 µM pAz Alexa488 and 7.2 µM pAz Alexa647 (PrF248+PrF358 VFT closure sensor) in acquisition buffer (20 mM Tris-HCl pH7.4, 118 mM NaCl, 1.2 mM KH2PO4, 1.2 mM MgSO4, 4.7 mM KCl, 1.8 mM CaCl2). After labeling, adherent cells were washed three times with DPBS w/o Ca^2+^ and Mg^2+^ (Thermo Fisher Scientific, France) for 5 min at RT each, followed by detachment of the cells in DPBS using a cell scraper. Cells were then collected by a 5 min centrifugation at 1,000 xg at RT, resuspended in cold lysis buffer (10 mM HEPES pH7.4, cOmplete™ protease inhibitor) on ice, incubated for 30 min and frozen at −80 °C. The following day, the lysis mixture was thawed in the fridge during 1 h, followed by 30 passages through a 200 µl pipette tip on ice. After two rounds of centrifugation at 500 xg, 4°C for 5 min each, the supernatant was centrifuged for 30 min, 4°C at 21,000 xg to collect the crude membranes. These were then overlaid once with acquisition buffer, recentrifuged for 5 min and flash frozen in liquid nitrogen after removal of the supernatant. Fractions were stored at -80 °C until solubilization of receptors.

Receptors were solubilized using 10 µl acquisition buffer containing 1% LMNG (w/v), 0.1% CHS Tris (w/v) per membrane fraction (corresponding to cells cultured in one well of a 6 well plate) for 15 min on ice. Subsequently, the solubilization mixture was centrifuged for 10 min at 4°C and 21,000 xg and the supernatant was mixed with 90 µl acquisition buffer containing 0.11% GDN (w/v). The diluted sample was then passed through a Zeba Spin Desalting Column (7 kDa cut-off, Thermo Fisher Scientific, France) equilibrated in acquisition buffer containing 0.005% LMNG (w/v), 0.0005% CHS Tris (w/v), 0.005% GDN (w/v). The eluate was then diluted 10 times in acquisition buffer and stored on ice in the dark.

### Functional integrity of solubilized receptors by LRET

The functional integrity of solubilized receptors was verified by LRET measurements after SNAP-tag labeling with SNAP-Lumi4-Tb and SNAP-green as described above. In brief, solubilized wildtype rat receptors and C1/C2 heterodimers were further diluted 2- and 4-times in acquisition buffer together with increasing concentrations of Glutamate or Glutamate + 10 µM BINA on white 384 well plates (polystyrene, flat-bottom, small volume, medium-binding, Greiner Bio-One SAS, France). Dose-response curves at different time points after storage of the samples at RT were obtained by LRET acquisitions on a Spark 20M (Tecan) through calculation of the ratio of the integrated sensitized acceptor emission at 535/25 nm at 100-500 µs over 1200-1600 µs after donor excitation at 340/20 nm.

### PIE-MFD smFRET setup

Single-molecule FRET experiments with pulsed inter- leaved excitation (PIE) – multiparameter fluorescence detection (MFD) were performed on a homebuilt confocal microscope ^37^ using the SPCM 9.85 software (B&H) as described previously^17^. Modifications are described in the following. A combination of 530/20 (530AF20, Omega Optical, Brattleboro, VT, USA) and 530/10 nm (FLH532-10, Thorlabs, Maisons-Laffitte, France) bandpass filters was used for Cy3B and Alexa546 excitation. A 488/10 (Z488/10 X, Chroma, Bellows Falls, VT, USA) bandpass filters was used for Alexa488 excitation. A 635/10 (FLH635-10, Thorlabs, Maisons-Laffitte, France) bandpass filter was used for “red” and Alexa647 excitation. The excitation power was 30 µW (prompt at 535 nm), 12 µW (delayed at 635 nm) for Cy3B / “red”; 50 µM (prompt at 535 nm), 15 µW (delayed at 635 nm) for Alexa546 / Alexa647; 25 µW (prompt at 488 nm), 12 µW (delayed at 635 nm) for Alexa488 / Alexa647 at the entrance into the microscope. Inside the microscope, the light was reflected by dichroic mirrors that match the excitation/emission wavelengths of the respective fluorophore combinations (Cy3B/Alexa546 with “red”/Alexa647: FF545/650- Di01, Semrock, Rochester, NY, USA and Alexa488 with Alexa647: FF500/646- Di01, Semrock, Rochester, NY, USA) and coupled into a 100 x, NA1.4 objective (Nikon, France). The following emission filters were used: Cy3B/Alexa546 parallel ET BP 585/65, perpendicular HQ 590/75 M (Chroma, Bellows Falls, VT, USA); Alexa488 parallel 535/50 BrightLine HC, perpendicular 530/43 BrightLine HC (Semrock, Rochester, NY, USA); red/Alexa647 parallel and perpendicular FF01-698/70-25 (Semrock, Rochester, NY, USA). Dual color emission was separated using FF649LP long pass filters (parallel and perpendicular, Semrock, Rochester, NY, USA) for Cy3B/Alexa546 with “red”/Alexa647 and AT608LP (parallel, Chroma, Bellows Falls, VT, USA) together with FF560LP (perpendicular, Semrock, Rochester, NY, USA).

### smFRET measurements

smFRET measurements were performed on SensoPlate 384 well plates (non-treated, Greiner Bio-One, France) passivated with 1 mg/ml bovine serum albumin (BSA) in acquisition buffer with 0.0025% LMNG (w/v), 0.00025% CHS Tris (w/v), 0.0025% GDN (w/v) for at least 1 h prior to sampling application. Samples containing 30-100 pM of labeled receptors were measured in acquisition buffer (20 mM Tris-HCl pH7.4, 118 mM NaCl, 1.2 mM KH2PO4, 1.2 mM MgSO4, 4.7 mM KCl, 1.8 mM CaCl2) with either 0.0025% LMNG (w/v), 0.00025% CHS Tris (w/v), 0.0025% GDN (w/v) for Cy3B/red or Alexa546/647 labeled constructs and 0.005% LMNG (w/v), 0.0005% CHS Tris (w/v), 0.005% GDN (w/v) for Alexa488/647 labeled constructs. Measurements at saturating ligand concentrations were performed at 10mM Glu, 100 µM LY379268, 100 µM LY341495, 1 mM DCG-IV either in the absence or presence of 1µM (Alexa488/647 constructs) or 10 µM BINA (Cy3B/red and Alexa546/647 constructs). The addition of BINA resulted in an increased fluorescent background when excited at a wavelength of 488 nm. We found that applying 1 µM kept the fluorescent background low enough to separate it from single molecule photon bursts of Alexa488-labeled receptors, while still promoting full VFT reorientation (Figure S6f). The heterotrimeric Gi1 complex was a kind gift from Sebastien Granier and Remy Sounier (IGF Montpellier, France) and was applied at 1 µM final concentration as previously described in^17^.

### smFRET data analysis

Data analysis was performed using the PAM 1.3 software package^38^. A single-molecule event was defined as a burst containing at least 40 photons with a maximum allowed interphoton time of 0.16 ms and a Lee-filter of 30. Apparent FRET efficiencies (EPR), accurate FRET efficiencies (E) and Stoichiometry (S) were calculated as described previously ^17^ following the recommendations made in ^39^ and ^40^.

Values for donor leakage α (fraction of the donor emission into the acceptor detection channel), direct excitation δ (fraction of the direct excitation of the acceptor by the donor-excitation laser), γ (correction factor that considers effective fluorescence quantum yields and detection efficiencies of the acceptor and donor) and β (correction factor that considers the relative laser excitation intensities) were α = 0.22, δ = 0.09 (SNAP- Cy3B/red), α = 0.1, δ = 0.13, γ = 0.7, β = 0.95 (248-Alexa546/647), α = 0.08, δ = 0.12, γ = 0.9, β = 1.2 (358- Alexa546/647) and α = 0.02, δ = 0.05, γ = 1.2, β = 0.75 (248+358-Alexa488/647).

To display FRET histograms, doubly-labeled molecules with an S = 0.3-0.7 (S = 0.35-0.75 for SNAP- Cy3B/red) were selected and normalized for the same number of molecules based on the macrotime of the experiment for individual biological replicates. In this way, average histograms with the mean FRET efficiency +/- SEM were obtained. Fitting was performed using Origin 6 (Microcal Software, Inc.) on average FRET histograms and for each individual data set separately to derive data shown in scatter plots. Histograms and plots were displayed using GraphPad PRISM 7.05.

### Additional software

The structures shown were generated using PYMOL 2.3.3. Figures were generated using Microsoft PowerPoint 2019 and INKSCAPE 0.92

## Authors contributions

NLG : designed research, designed and performed molecular biology, chemical synthesis, UAA screening, functional test of receptors, LRET titrations and microscopy, analyzed and interpreted data; prepared figures

RBQ : designed and performed protein expression, solubilization, labeling and data acquisition for smFRET; analyzed and interpreted data; prepared figures; wrote the manuscript

HL : designed and performed molecular biology, UAA screening, functional test of receptors, LRET experiments

TCM : designed, supervised and analyzed LRET microscopy experiments, critically revised the manuscript XR : performed molecular modeling to select positions for UAA incorporation

SS : technical help in the organic synthesis of compounds LL : design organic synthesis

ET, JL : supervised research and interpreted the data

JPP, PR, EM : Designed and supervised research, interpreted data, acquired funds, wrote the manuscript

## Acknowledgments

Our research is supported by grants from the Agence Nationale pour la Recherche (ANR-09-PIRI-0011 to PR, ANR 17-CE09-0026-02 to EM and ANR 18-CE11-0004-02 to EM and JPP) ; the Fondation Recherche Médicale (DEQ20170336747 to JPP). P. R. and J.-P. P. were by supported by the Institut National de la Santé et de la Recherche Médicale (INSERM ; International Research Program « Brain Signal ») and the Franco-Chinese Joint Scientific and Technological Commission (CoMix) from the French Embassy in China. H.L. was supported by Sino-French Cai Yuanpei program. J.L. was supported by the National Natural Science Foundation of China (NSFC) (grant numbers 31130028 and 31225011 to J.L) ; the Ministry of Science and Technology (grant number 2012CB518000 to J.L.) ; the Program for Introducing Talents of Discipline to the Universities of the Ministry of Education (grant number B08029 to J.L.). X. R. by a FEBS long term fellowship and by a Agència de Gestió d’Ajuts Universitaris i de Recerca (AGAUR) BP post-doctoral fellowship. The CBS belongs to the France-BioImaging national infrastructure supported by the French National Research Agency (ANR-10-INBS-04, “Investments for the future”) and is supported by the GIS “IBiSA: Infrastructures en Biologie Santé et Agronomie”. We thank the members of the IBM team (CBS, Montpellier) for fruitful discussions; the Arpège platform (IGF, Montpellier) for providing facilities and technical support, Perkin Elmer Cisbio for providing reagents. We thank Thierry Durroux (IGF, Montpellier) for setting-up the LRET microscope and imaging experiments, Emmanuel Bourrier (Cisbio) for his help with dye synthesis, Thomas P Sakmar (Rockfellar University, NY, USA), Pierre Paoletti (IBENS, ENS, Paris, France), Irene Coin and Robert Serfling (Universität Leipzig, Germany) for providing plasmids for ncAA incorporation.

