## Supplementary Figures for "Dissecting conformational rearrangements and allosteric modulation in metabotropic glutamate receptor activation"

a

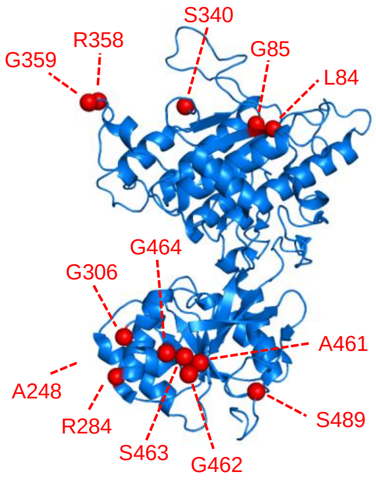


b

| Position | VFT Lobe | Yield (% WT) | Functional (IP-One) |
|---|---|---|---|
| 84 | Upper lobe | 21.2 +/-1.1 | Yes |
| 85 | Upper lobe | 23.4 +/-0.7 | No |
| 340 | Upper lobe | 12.3 +/-0.5 | Yes |
| 358 | Upper lobe | 29.8 +/-0.3 | Yes |
| 359 | Upper lobe | 32.6 +/-4.7 | Yes |
| 248 | Lower lobe | 35.9 +/-3.2 | Yes |
| 284 | Lower lobe | 29.5 +/-2.2 | Yes |
| 306 | Lower lobe | 25.5 +/-4.5 | Yes |
| 461 | Lower lobe | no | NA |
| 462 | Lower lobe | no | NA |
| 463 | Lower lobe | no | NA |
| 464 | Lower lobe | no | NA |
| 489 | Lower lobe | no | NA |
| 248/84 | Both | 5.8 +/-0.7 | Yes |
| 248/85 | Both | 11.8 +/-1.6 | No |
| 248/340 | Both | 15.7 +/-3.5 | No |
| 248/358 | Both | 36.1 +/-1 | Yes |
| 248/359 | Both | 17.1 +/-2 | Yes |

**Supplementary Figure 1: Screening of positions for incorporation of ncAAs.** a) Structure of a single VFT (PDB 7EPA) with positions screened for PrF incorporation highlighted in red. b) Summary of screening results showing the positions substituted by PrF, their location within the VFT, their incorporation efficiency relative to expression of the wildtype receptor and their functionality. Expression relative to the wildtype was determined by time-resolved FRET measurements of N-terminally labeled SNAP-tags with BG-Lumi4-Tb. Functionality was evaluated by the increase in inositol phosphate accumulation in response to activation with saturating LY379268 as compared to no activation in the absence of ligand using the IPone assay (Cisbio, France). Data represent the mean +/-SEM of duplicate analysis. NA: not applicable.

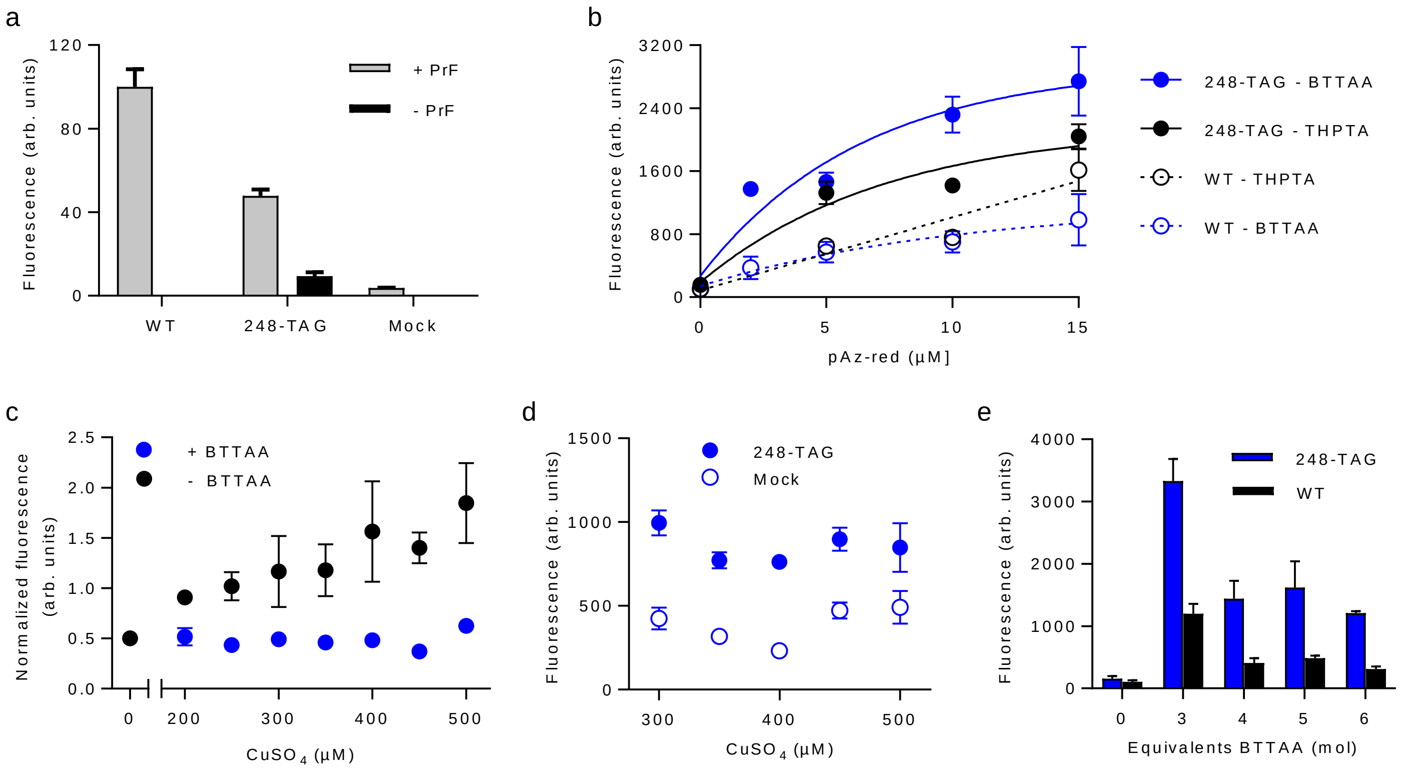


**Supplementary Figure 2: Optimization of live cell compatible click chemistry conditions.** a) PrF-dependent suppression of premature TAG at position 248 (248-TAG) leads to expression of full-length SNAP-mGlu2-PrF248 as shown by cell surface specific SNAP-labeling with BG-Lumi4-Tb. b) Labeling efficiency of CuAAC reaction using BTTAA (blue, 360 µM $CuSO_4$, 6 eq. BTTAA) or THPTA (black, 300 µM $CuSO_4$, 6.6 eq. THPTA). Labeling was performed on cells expressing SNAP-mGlu2-PrF248 or wildtype receptor with 15 µM of pAz-red for 25 min at 37°C. c) Cytotoxicity test in response to increasing concentrations of $CuSO_4$ in the absence (black) or presence of 6 eq. BTTAA. Cytotoxicity was determined by the increase in fluorescence as a result of propidium iodide staining, normalized by staining with Hoechst 33342. d) Influence of $CuSO_4$ concentration on labeling efficiency using 6 eq. BTTAA and 15 µM pAz-red. e) Influence of molar ratio between $CuSO_4$ (360 µM) and BTTAA on labeling efficiency (248-TAG) and specificity (WT) using pAz-red. All data are represented as the mean +/- SEM of triplicates.

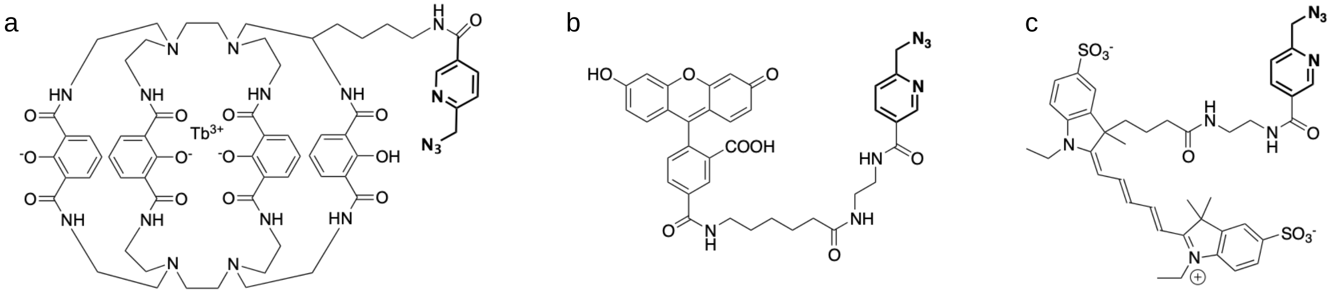


| | | |
|---|---|---|
| pAz-Lumi4-Tb | pAz-green | pAz-red |
| $C_{63}H_{74}N_{17}O_{13}Tb$ | $C_{36}H_{33}N_7O_8$ | $C_{41}H_{49}N_8O_8S_2$ |
| MW 1435.4906 Da | MW 691.1391 Da | MW 845.3109 Da |

**Supplementary Figure 3: Structures of Lumi4-Tb (a), "green" (b) and "red" (c) Picolyl-azide (pAz) dye derivatives.**

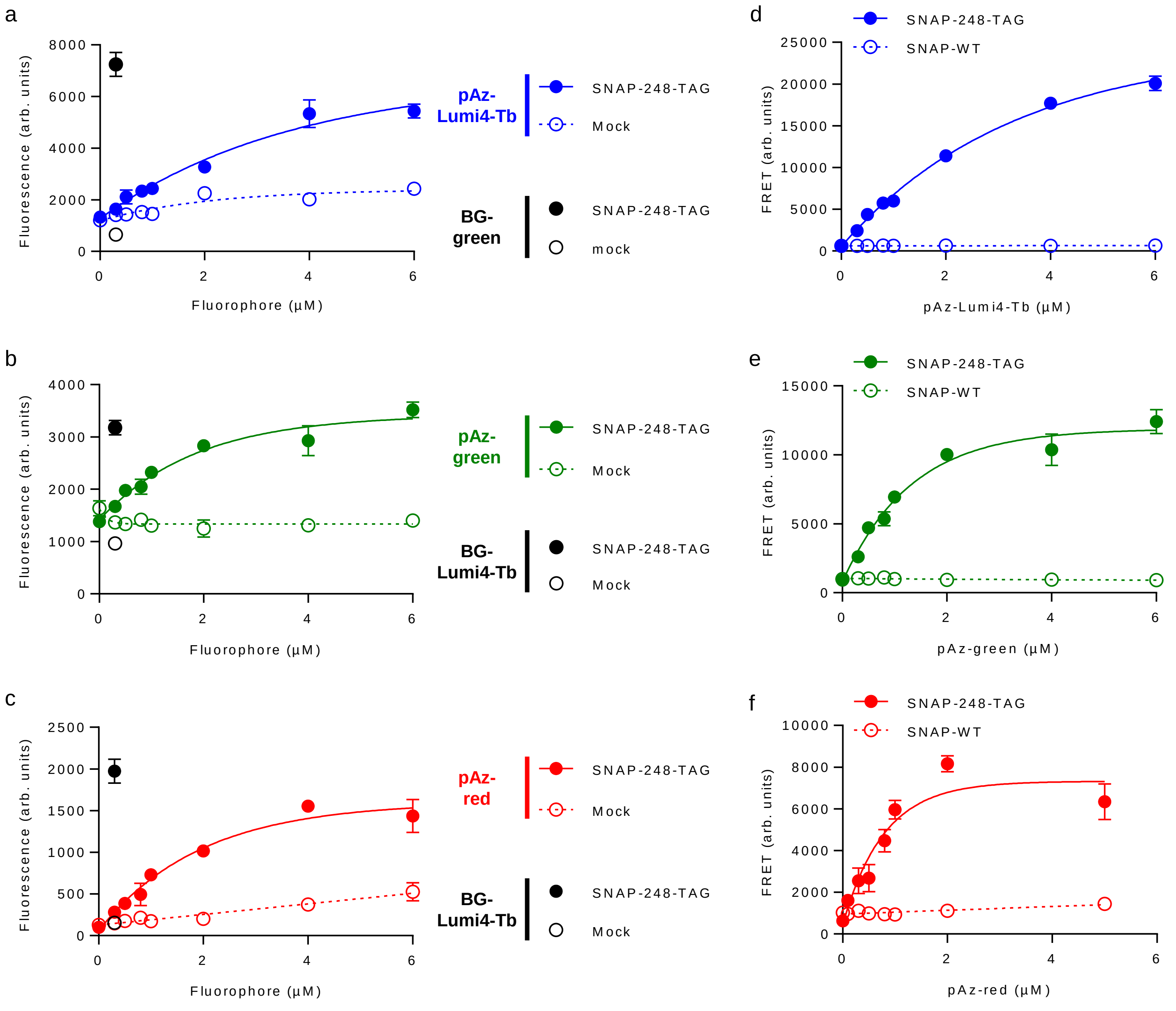


**Supplementary Figure 4: Optimization of labeling with pAz dyes for LRET measurements.** a-c) Labeling efficiency of SNAP-mGlu2-PrF248 (SNAP-248-TAG, solid line) and specificity compared to cells transfected with empty vector (Mock, dashed line) using CuAAC at increasing concentrations of indicated pAzF dyes. Maximal and non-specific labeling of SNAP-tag with corresponding BG-dye derivatives is also shown. d-f) Increase in FRET signal in response to labeling with increasing concentrations of pAz dyes. The FRET signal is given as sensitized emission of acceptor after Lumi4-Tb donor excitation by LRET. Labeling was performed with BG-green (d) or BG-Lumi4-Tb (e-f) followed by CuAAC with pAz-Lumi4-Tb (d), pAz-green (e) or pAz-red (f). All data are represented as the mean +/- SEM of triplicates.

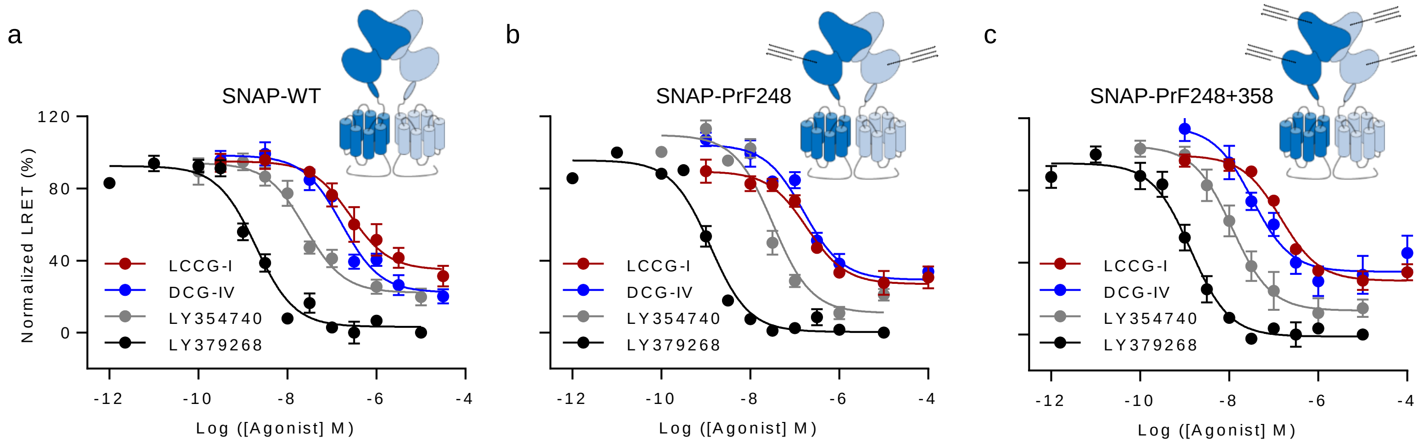


**Supplementary Figure 5: Functional characterization of wildtype receptor and variants with incorporated PrF.** Dose-response curve, normalized to LY379268, were obtained by ligand titration of cells expressing wildtype receptors (a, SNAP-WT) and receptors with PrF incorporated at position 248 (b, SNAP-PrF248) or 248 and 358 (c, SNAP-PrF248+358). Receptor activation is presented as a decrease in LRET signal due to accumulation of inositol phosphate (IPone assay). Data represent the mean +/- SEM of 3-12 replicates.

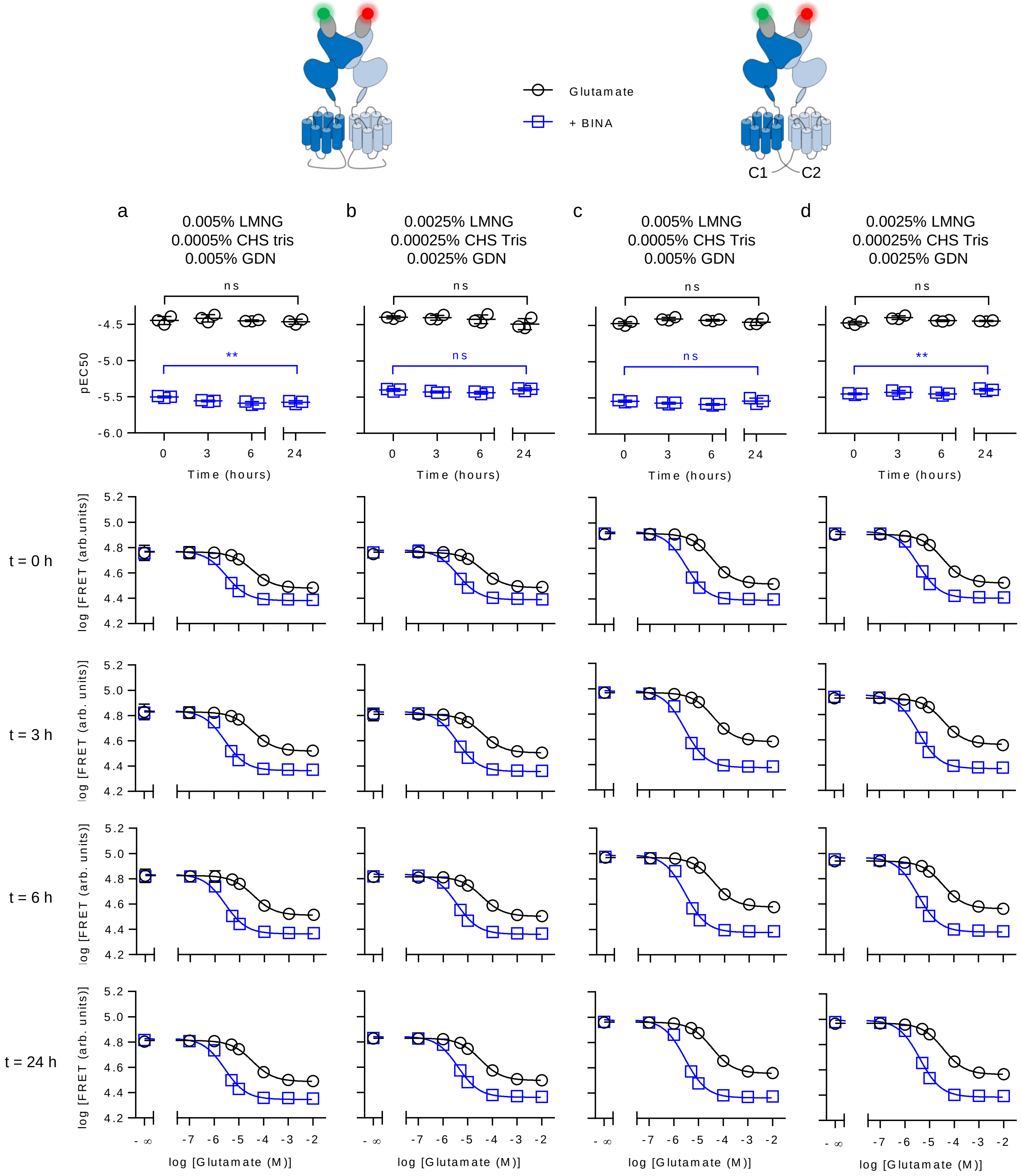


**Supplementary Figure 6: Evaluation of functional integrity of receptors over time after solubilization using LRET.** Wildtype receptor homodimers (a-b) or heterodimers containing the engineered C-terminal C1 and C2 $GABA_B$ quality control system (c-d) were labeled with BG-Lumi4-Tb donor and BG-green acceptor via N-terminal SNAP-tags and solubilized from membrane fractions using 1% LMNG + 0.1% CHSTris. Subsequently, supernatants were diluted in the presence of GDN to reach final concentrations of either 0.005% LMNG + 0.0005% CHS Tris + 0.005% GDN (a and c) or 0.0025% LMNG + 0.00025% CHS Tris + 0.0025% GDN (b and d) in acquisition buffer and titrated with glutamate alone (black) or glutamate + 10 µM BINA (blue). Functional integrity over time at room temperature was evaluated based on the changes in glutamate and glutamate + BINA pEC50 values (top row) obtained from dose-response curves performed on three independent biological replicates analyzed in one experiment and are presented together with the mean +/- SEM. Statistical differences were determined using two-sided, unpaired t-tests and are given $^{**}p \leq 0.01$, ns > 0.05. The average dose-response curves were obtained by fitting the mean +/- SEM at t = 0, 3, 6 and 24h of sample storage at room temperature (from top to bottom) are shown below the corresponding pEC50 plots with.

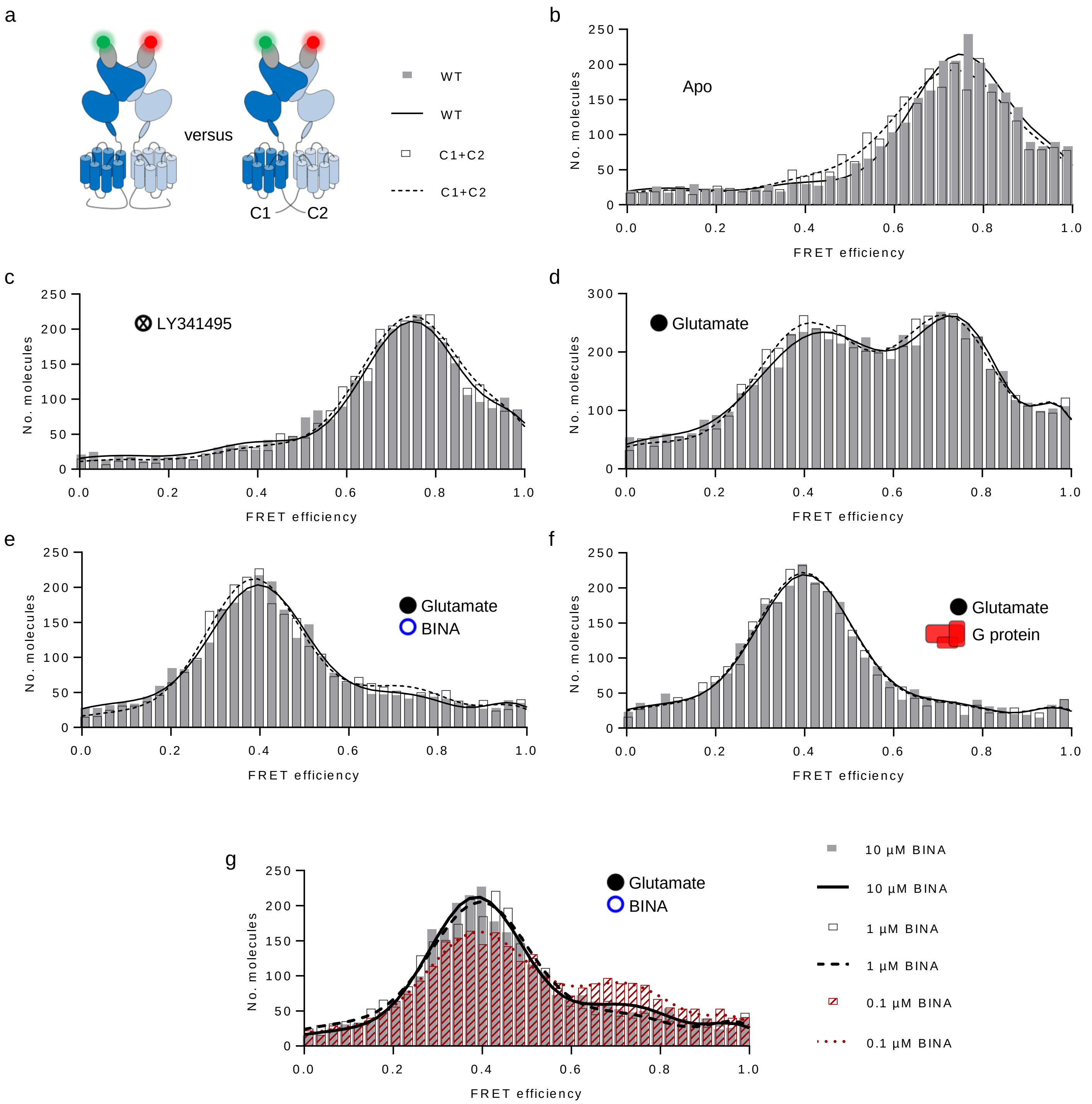


**Supplementary Figure 7: Comparison of ligand-induced VFT reorientation between wildtype and C-terminally modified mGlu2.** Receptor constructs were labeled with BG-Cy3b donor and BG-D2 acceptor on N-terminal SNAP-tags, solubilized and VFT reorientation measured by smFRET at different ligand conditions. a) Shown are representative FRET histograms of wildtype receptor (WT, gray bars, global fit given as black solid line) superimposed on histograms obtained for C-terminally modified receptor (C1+C2, white bars, global fit given as black dashed line) in the absence of ligand (b, Apo), the presence of saturating LY341495 (c), glutamate (d), glutamate + BINA (e) and glutamate + heterotrimeric $G_i$ protein (f). g) Superimposition of representative FRET histograms obtained for C-terminally modified SNAP-mGlu2, labeled as above, in the presence saturating glutamate together with 0.1 µM BINA (red bars with global fit given as red dashed line), 1 µM BINA (white bars with global fit given as black dashed line) and 10 µM BINA (gray bars with global fit given as solid black line). All histograms were obtained by plotting a fixed number of doubly labeled molecules (S = 0.35-0.75, a-b and d-f = 3000+/-2 molecules, c = 6000+/-2 molecules) versus the FRET efficiency given as $E_{PR}$ (corrected for direct excitation and crosstalk as specified in Methods).

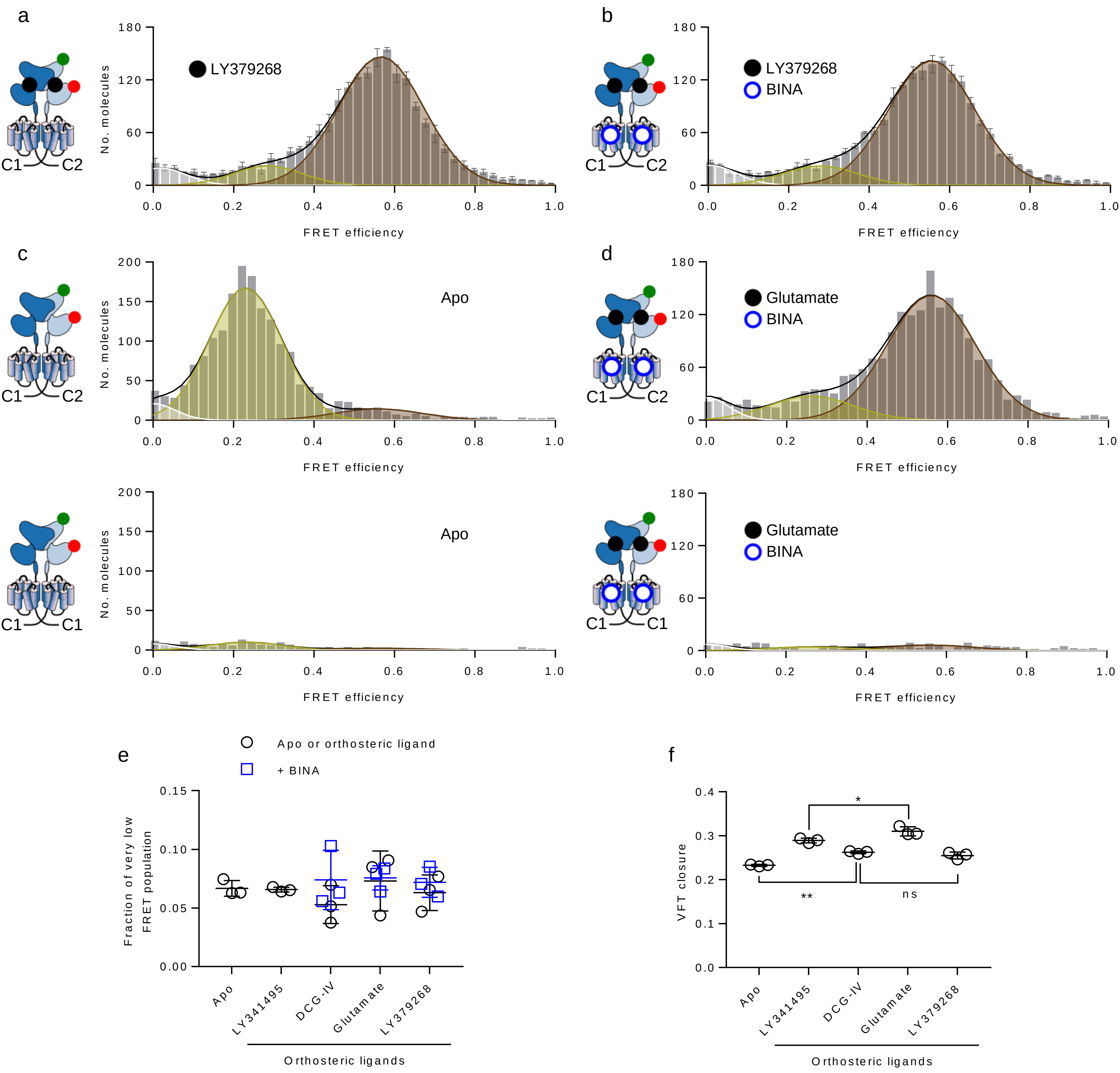


**Supplementary Figure 8: Supplementary data for VFT closure sensor.** FRET histograms of VFT closure sensor ($U_{358}$-$L_{248}$) in the presence of saturating LY379268 (a) and LY379268 + 1 µM BINA (b). FRET histograms show the accurate FRET efficiency as the mean +/- SEM of three independent biological replicates each normalized to 2000 events in the DA population (S = 0.3-0.7). Histograms display the fitting with 3 gaussians (white = very low FRET, yellow = low FRET, red = high FRET) together with the global fit (black). c-d) Representative FRET histograms of samples obtained by expression of 248TAG+358TAG-C1 in the presence (top) or absence (bottom) of WT-C2. Top and bottom histograms in the absence (c) or presence of glutamate + 1 µM BINA (d) were each plotted for the same measurement macrotime and corrected for the applied sample volume using samples that were prepared in the same manner in parallel (c: 8955 s at 16 µl = 4776 s at 30 µl, d: 11620 s at 13 µl = 5035 s at 30 µl) to evaluate the degree of signal obtained from leakage of the 248+358-C1 sensor to the cell surface in the absence of WT-C2. e) Representation of the fraction of population of the very low state in response to different ligand conditions given as the number of molecules found in the respective population over the sum of molecules in all three populations. Black circles show the Apo condition and orthosteric ligands, while blue squares are given for agonists in the presence of 1 µM BINA. f) Determination of the mean FRET efficiency of the low FRET (yellow) population (VFT closure) for the different ligand conditions shown in Figure 3b-e and S8a. Data in e-f are given with the mean +/- SEM of three independent biological replicates. f) . Statistical differences were determined using a one-way ANOVA with Sidak multiple comparisons test and are given as ****p ≤ 0.0001, ***p ≤ 0.001, **p ≤ 0.01, *p ≤ 0.05, ns > 0.05.

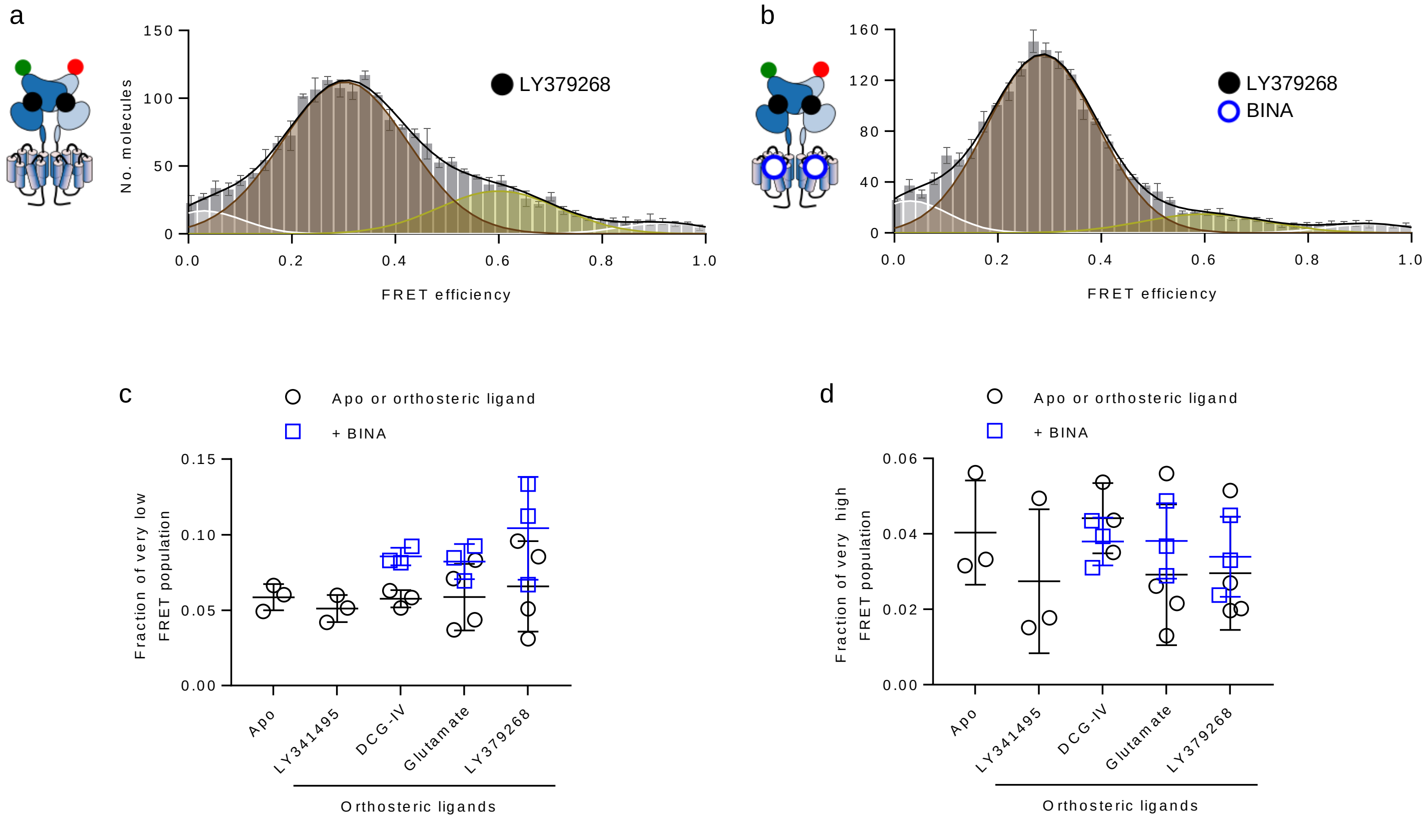


**Supplementary Figure 9: Supplementary data for upper lobe FRET sensor.** a-b) FRET histograms of upper lobe sensor ($U_{358}$-$U_{358}$) in the presence of a saturating concentration of the synthetic full agonist LY379268 alone (a) and in the presence of 10 µM BINA (b).Histograms show the accurate FRET efficiency as the mean +/- SEM of three independent biological replicates each normalized to 2000 events in the DA population (S = 0.3-0.7). Histograms display the fitting with 4 gaussians (yellow = LF, red = HF, white = VLF and VHF) together with the global fit (black). c-d) Representation of the fraction of population of the very low (c) and very high (d) FRET states in response to different ligand conditions given as the number of molecules found in the respective population over the sum of molecules in all four populations. Black circles show the Apo condition and orthosteric ligands while blue squares are given for agonists in the presence of 10 µM BINA.

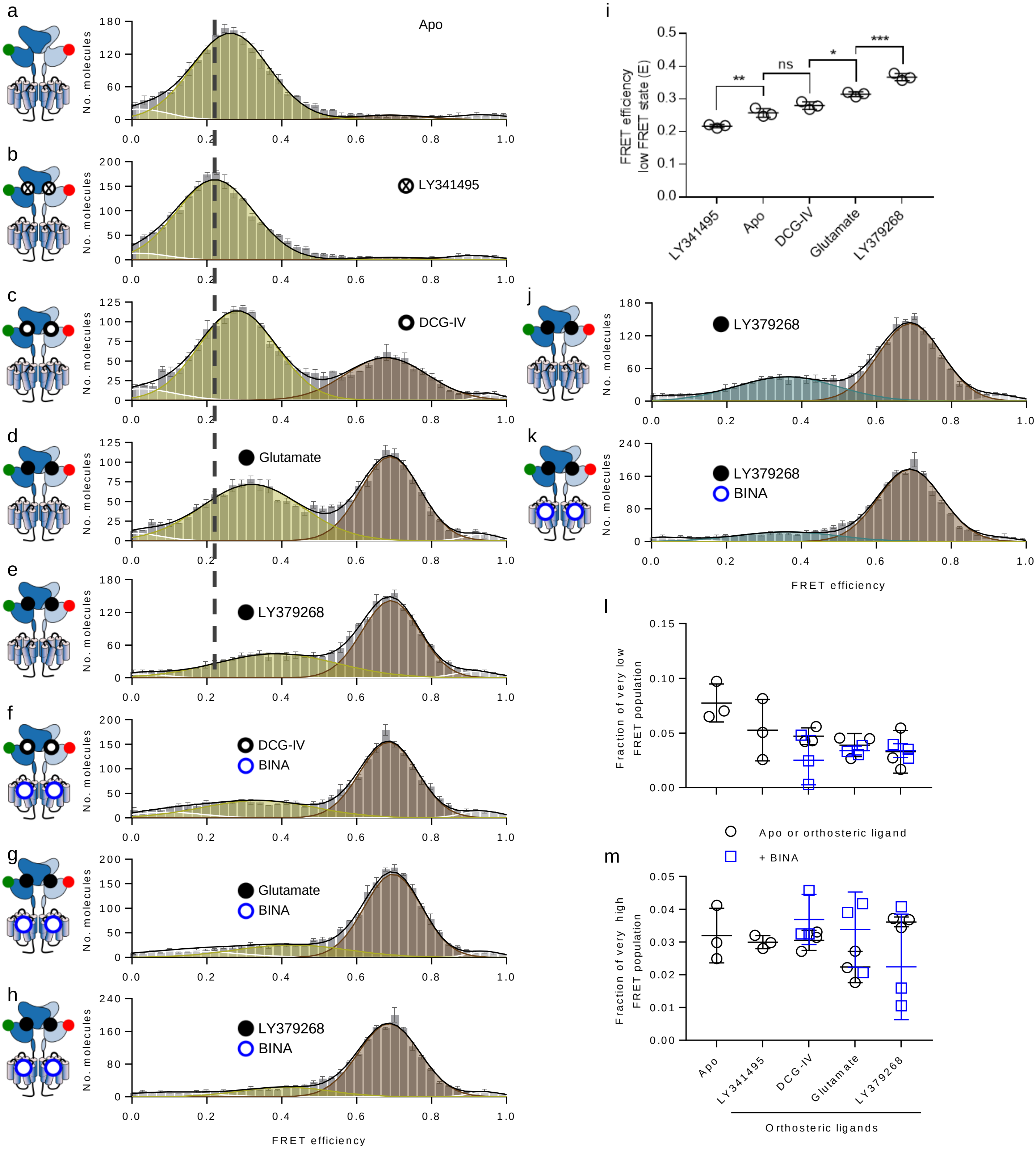


**Supplementary Figure 10: Supplementary data for lower lobe FRET sensor.** a-h) FRET histograms of lower lobe sensor ($L_{248}$-$L_{248}$) in the absence of ligand (Apo, a) and in the presence of saturating LY341495 (b), DCG-IV (c), Glutamate (d), LY379268 (e), DCG-IV + BINA (f), Glutamate + BINA (g) and LY379268 + BINA (h). Histograms display the same data as in Figure 5b-g but fitted with 4 gaussians (white = very low FRET, yellow = low FRET, red = high FRET, white = very high FRET) together with the global fit (black). i) Mean FRET efficiency of low FRET state (yellow) in a-e. Statistical differences were determined using a one-way ANOVA with Sidak multiple comparisons test and are given as ****$p \leq 0.0001$, ***$p \leq 0.001$, **$p \leq 0.01$, *$p \leq 0.05$, ns $> 0.05$. j-k) FRET histograms in the presence of a saturating concentration of the synthetic full agonist LY379268 alone (g) and in the presence of 10 µM BINA (h). Histograms display the fitting with 5 gaussians (yellow = low FRET1, blue = low FRET2, red = high FRET, white = very low FRET and very high FRET) together with the global fit (black). FRET histograms in a-h and j-k show the accurate FRET efficiency as the mean +/- SEM of three independent biological replicates each normalized to 2000 events in the DA population (S = 0.3-0.7). l-m) Representation of the fraction of population of the very low (l) and very high (m) FRET states in response to different ligands, given as the number of molecules found in the respective population over the sum of molecules in all four populations shown in a-h. Black circles show the Apo condition and orthosteric ligands [illegible]
